# Niche signals regulate continuous transcriptional states in hematopoietic stem cells

**DOI:** 10.1101/2021.03.09.430613

**Authors:** Eva M Fast, Audrey Sporrij, Margot Manning, Edroaldo Lummertz da Rocha, Song Yang, Yi Zhou, Jimin Guo, Ninib Baryawno, Nikolaos Barkas, David T Scadden, Fernando Camargo, Leonard I Zon

## Abstract

Hematopoietic stem cells (HSCs) must ensure adequate blood cell production following distinct external stressors. A comprehensive understanding of *in vivo* heterogeneity and specificity of HSC responses to external stimuli is currently lacking. We performed single-cell RNA sequencing (scRNA-Seq) on functionally validated mouse HSCs and LSK (Lin-, c-Kit+, Sca1+) progenitors after *in vivo* perturbation of niche signals interferon, granulocyte-colony stimulating factor (G-CSF), and prostaglandin. We identified six HSC states that are characterized by enrichment but not exclusive expression of marker genes. Niche perturbations induce novel and rapid transitions between these HSC states. Differential expression analysis within each state revealed HSC- and LSK-specific molecular signatures for each perturbation. Chromatin analysis of unperturbed HSCs and LSKs by scATAC-Seq revealed HSC-specific, cell intrinsic predispositions to niche signals. We compiled a comprehensive resource of HSC- and progenitor-specific chromatin and transcriptional features that represent important determinants of regenerative potential during stress hematopoiesis.

## Introduction

Stem cell therapy holds promises for numerous indications, including blood diseases, autoimmune diseases, neurodegeneration and cancer (Blau and Daley, 2019). Despite being used in the clinic for over 30 years, HSC transplants remain a highly risky procedure. To better understand HSC regeneration, recent efforts have used scRNA-Seq to discover novel markers to further enrich for functional HSCs (Chen et al., 2016, Cabezas-Wallscheid et al., 2017, Wilson et al., 2015, Rodriguez-Fraticelli et al., 2020). Yet no consensus exists on the optimal marker combination to obtain the most purified HSCs in part because extensive functional heterogeneity within HSCs makes experimental evaluation challenging (Haas et al., 2018). Both intrinsic and extrinsic factors have been implicated in regulating HSC function (Zon, 2008, Morrison et al., 1996). The stem cell niche forms an important extrinsic regulator of HSCs as it anchors stem cells and maintains the balance between self-renewal and differentiation (Morrison and Spradling, 2008, Morrison and Scadden, 2014). Release of soluble signals from the niche such as interferons, prostaglandins, and growth factors, including SCF and G-CSF, have been shown to influence HSC function during homeostasis and upon injury (Pinho and Frenette, 2019, Pietras et al., 2016, Zhao et al., 2014, Morales-Mantilla and King, 2018). While known to be affected by a wide variety of extracellular signals, little is known about the heterogeneity and specificity of HSC responses to these external stimuli, nor is it understood how differential responses relate to functional diversity of HSCs. HSCs are also regulated cell intrinsically (Zon, 2008, Morrison et al., 1996). Chromatin state is a crucial determinant of cell identity and behavior (Klemm et al., 2019). Hematopoietic differentiation is a prime example of how cell fate changes associate with massive remodeling of the epigenetic landscape (Avgustinova and Benitah, 2016). Despite the current knowledge on regulators of HSC fate, few studies have assessed chromatin states in purified, *in vivo* derived HSC populations (Yu et al., 2017, Lara-Astiaso et al., 2014) due to technical limitations such as cell numbers. Recent advancements in single cell chromatin accessibility sequencing (scATAC-Seq) provides a methodological framework for studying the diversity and uniqueness of HSC chromatin features at homeostasis and upon external stimulation (Buenrostro et al., 2018, Lareau et al., 2019). Here, we performed comprehensive scRNA-Seq and scATAC-Seq profiling on functionally validated mouse HSCs and examined *in vivo* transcriptional responses to extracellular stimulation, mimicking signals from the stem cell niche. We found that unperturbed HSCs exist in distinct transcriptional states. Niche signals can alter the cell distribution between HSC states to varying degrees depending on the stimulant as well as induce specific changes within cell states. Comparison of HSCs to multipotent LSK (Lin-, c-Kit+, Sca1+) progenitors allowed us to determine the specificity of transcriptional responses in HSCs. Finally, analysis of native HSC chromatin states revealed cell intrinsic heterogeneity that may prime HSC subpopulations for particular transcriptional responses following exposure to certain signals. The data is provided as a resource to the broader research community via an easily accessible web interactive application (https://mouse-hsc.cells.ucsc.edu). Overall, this work provides the first comprehensive description of the single cell transcriptomic and epigenetic landscape of HSCs and multipotent LSK progenitors *in vivo*.

## Results

### *In vivo* stimulation of functionally validated HSCs and multipotent progenitors (MPPs) for transcriptomic and epigenetic profiling

To investigate transcriptional responses to external signals, we profiled HSCs after four distinct *in vivo* niche perturbations (Fig 1A). Male and female mice were treated with one of three activators, 16,16-dimethyl Prostaglandin E_2_ (dmPGE_2_), Poly(I:C), or G-CSF for two hours or administered the Cox1/2 inhibitor Indomethacin (here abbreviated to ‘Indo’) for one week to deplete endogenous prostaglandins (see Methods). After the respective drug treatments, HSC and MPP populations comprising the entire LSK compartment were isolated via FACS (Sup Fig 1A). Through limit dilutions transplantation assay (LDTA) and ELDA analysis (Hu and Smyth, 2009), we determined HSC purity to be 1 in 8 (Sup Fig 1B-1D). This confirmed that our isolation and purification procedure allows for the profiling of functional, highly purified HSCs. Cell cycle analysis further verified that HSCs were mostly quiescent, in contrast to other MPP populations (Sup Fig 1E)(Cabezas-Wallscheid et al., 2014). Phenotypic marker composition within LSK cells remained largely consistent between different stimulations (Sup Fig 1F). An exception was the reduction of cells within the HSC compartment following dmPGE_2_ treatment, decreasing from 1.9% in control to 0.85% of LSK cells (p-value = 6.4*10^-4^, by differential proportion analysis (DPA)(Farbehi et al., 2019)). To account for a potential phenotypic shift in HSC surface marker expression due to CD34 externalization, which would move functional HSCs to the MPP1 population, we compared the contribution of the later by scRNA-Seq defined ‘stem cell state’ in HSCs and MPP1s. We found no increase in the ‘stem cell’ population in dmPGE_2_ treated MPP1s, compared to the control (Sup Fig 2G). After cell sorting, we subjected a total of 46,344 cells to scRNA-Seq using the 10x Genomics platform (see Methods). We obtained an average of 37,121 (SD = 14,308) reads per cell and 2,994 (SD = 480) genes per cell (Sup Table 1), indicative of a rich dataset that contained functionally validated HSCs.

**Figure 1:**
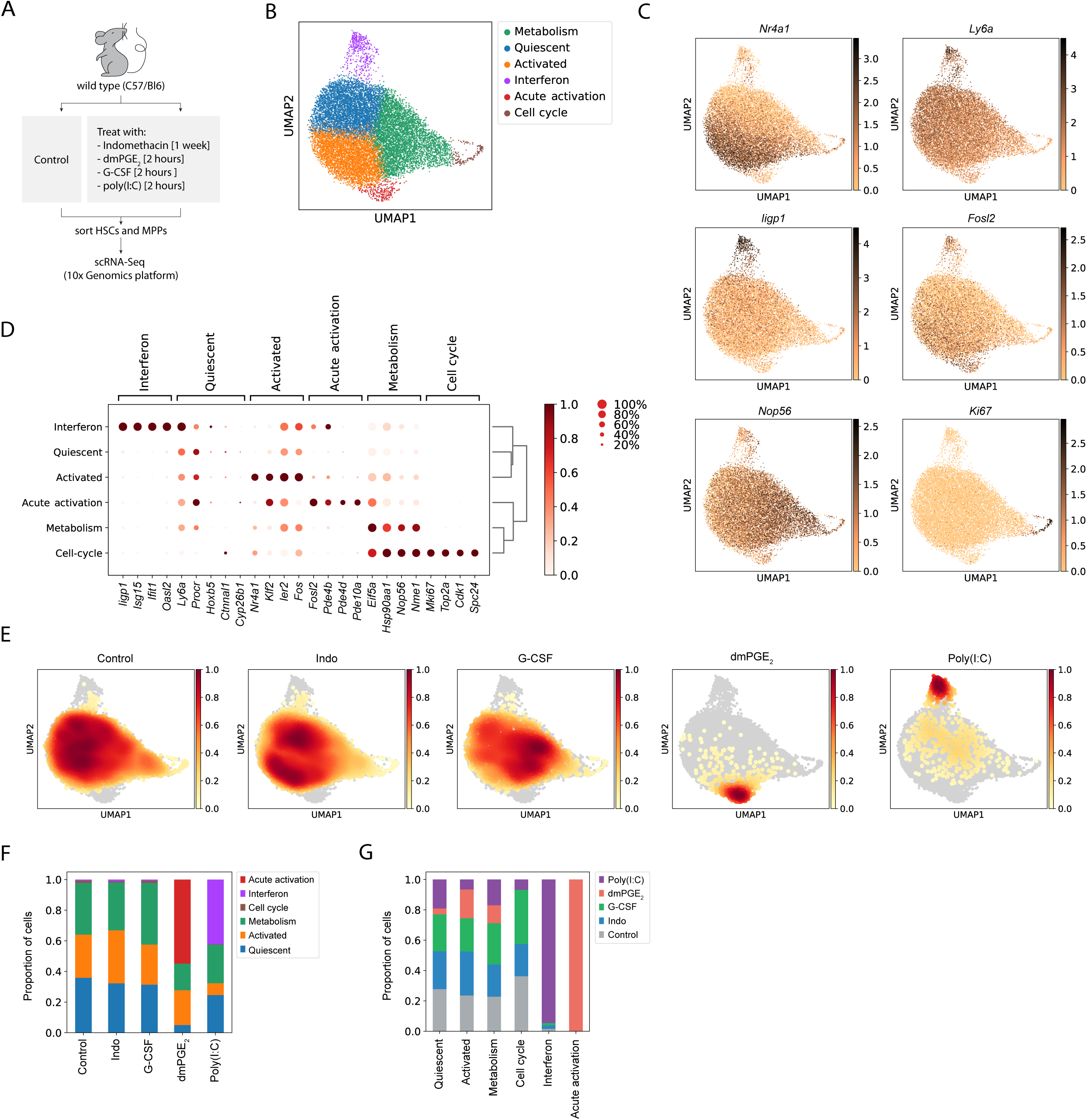
HSCs are transcriptionally heterogeneous and niche perturbations rapidly shift cells into different states. (A) Schematic of stimulant treatment before HSC and MPP isolation. (B) UMAP plot of HSC clusters (n = 15 355 cells) (C) UMAP plot with expression of representative genes for each cluster. (D) Dot plot of enriched genes for each HSC cluster (scaled expression across columns). (E) UMAP density graphs of HSC distribution upon each niche stimulant (F) Proportion of HSCs within clusters for each niche perturbation. (G) Proportion of HSCs from each niche perturbation within a cluster normalized for total cell number per treatment.

**Figure 2:**
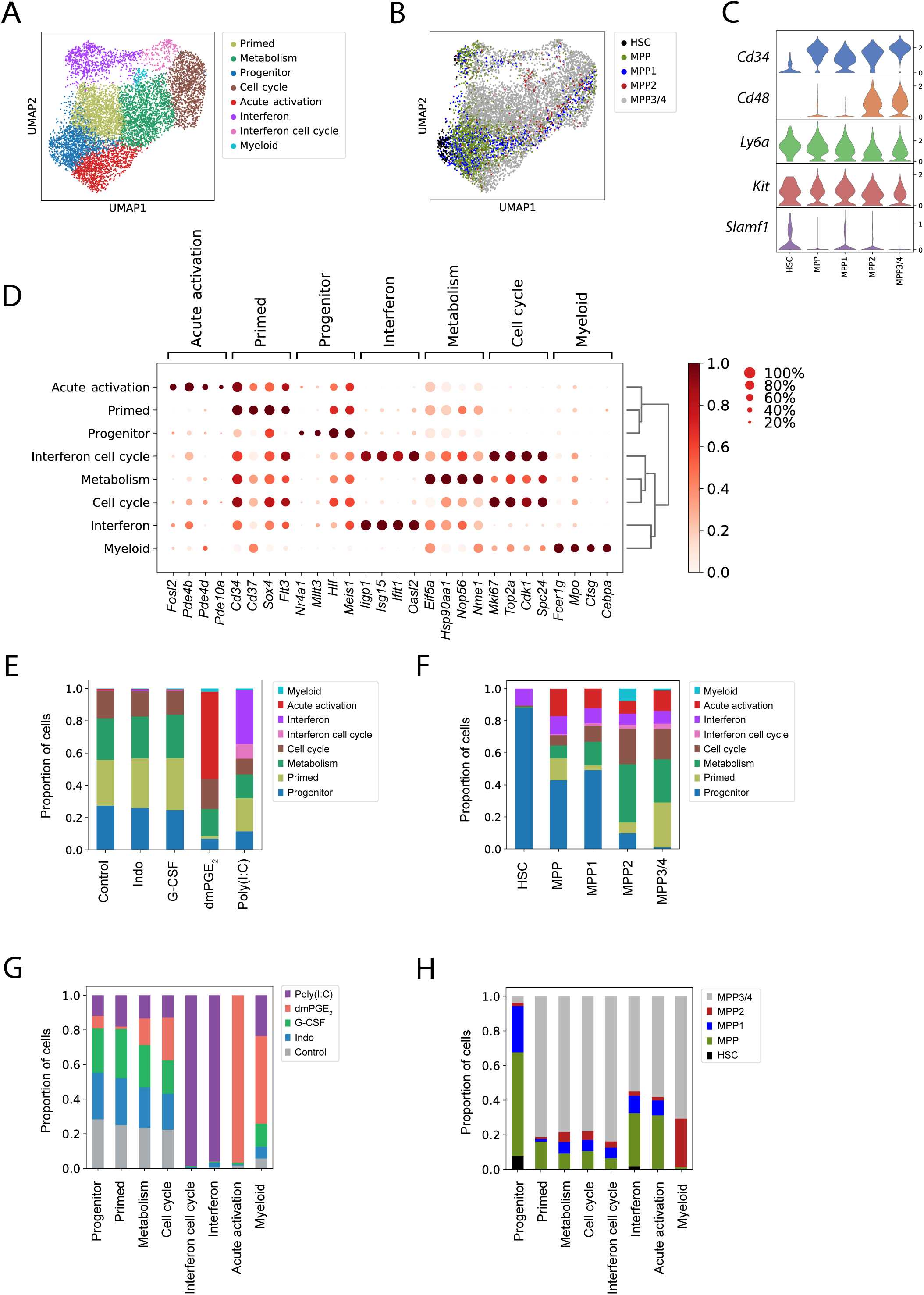
MPP surface marker validated LSK clusters respond similarly to drug treatments like HSCs. (A) UMAP plot of LSK clustering (n = 8191 cells). (B) UMAP plot of surface receptor phenotypes in LSK cells. (C) Stacked violin plots of gene expression for surface markers within MPPs and HSCs. (D) Dot plot of enriched genes for each LSK cluster (scaled expression across columns). (E) Proportion of LSK cells within clusters for each niche perturbation. (F) Proportion of LSK cells belonging to different clusters for each surface phenotype. (G) Proportion of LSK cells from each niche perturbation within a cluster normalized for total cell number per treatment. (H) Proportion of surface phenotypes within each LSK cluster.

### Niche signals induce rapid transitions between transcriptional HSC states

To determine how niche stimulants affect HSCs, we analyzed a combination of control and treated cells. We applied a standard scRNA-Seq pipeline to filter and normalize UMI reads (see Methods). Separate analysis of male and female HSCs revealed little sexual dimorphism during both steady state and following activation (Sup Fig 3, Sup Table 2 and 3). We therefore regressed out any sex specific effects from further downstream analyses (see Methods). In the aggregated dataset, we detected a total of six clusters (Fig 1B). To ensure optimal choice of clustering hyper parameters, we used a data driven approach (Silhouette Coefficient and Davies–Bouldin index) that was validated by comparison of two independent biological scRNA-Seq replicates of control HSCs sorted from different mouse strains (see Methods, Sup Fig 2A-2D, Sup Table 4). The absence of clear separation into highly distinct clusters in UMAP space (Fig 1B), together with fact that most marker genes were not exclusively expressed but rather enriched in a given cluster (Fig 1C), suggests that the HSC clusters represent transcriptional states with continuous transitions as opposed to discrete subtypes of HSCs. Reactome pathway enrichment analysis (Sup Fig 2E) in combination with manual curation of enriched genes (Fig1D, Sup Table 4) allowed for assignment of labels to each state/cluster. Three HSC clusters equally made up 98% of control cells (Fig 1F) and two minor clusters comprised 2% of control HSCs. Consistent with the FACS results (Sup Fig 1E), the proportion of HSCs residing in a ‘Cell cycle’ cluster marked by genes such as *Ki67* was very low (1%, Fig1C and 1F). A prominent HSC subpopulation contained cells that were defined by various immediate early genes (IEGs including *Nr4a1*, *Ier2* and *Fos* (Fig 1B – D). We therefore named this cluster ‘Activated’. We eliminated the possibility that the ‘Activated’ cluster arose due to an unspecific artifact of the cell isolation procedure since LSKs did not have an ‘Activated’ cluster and the proportion of *Nr4a1* expressing cells was much smaller (Fig 2D and Sup Fig 4B). HSCs have been tightly linked to dormancy and quiescent states (Foudi et al., 2009, Wilson et al., 2008, Qiu et al., 2014). The cluster adjacent to the ‘Activated’ state was termed ‘Quiescent’ because cells within this population showed the highest expression levels of previously described HSC markers (Cabezas-Wallscheid et al., 2017, Chen et al., 2016, Wilson et al., 2015, Acar et al., 2015, Gazit et al., 2014, Balazs et al., 2006) (Sup Fig 2F). Furthermore, ‘Quiescent’ HSCs did not express IEGs (Fig 1C) and were located most distal to the ‘Cell cycle’ cluster in UMAP space (Fig 1B-D). The ‘Metabolism’ cluster was comprised of the most metabolically active HSCs and showed enrichment of transcripts involved in translation initiation (*Eif5a, Eif4a1*), nucleotide metabolism (*Nme1, Dctpp1*), ribosome assembly (*Ncl, Nop56, Nop10, Npm1*), protein chaperones (*Hsp90, Hsp60*) and was located adjacent to the ‘Cell cycle’ cluster (Fig 1B-D). We next evaluated whether treatment with niche stimulants shifts the distribution of cells between clusters (Fig 1E). Interferons induced by poly(I:C) treatment increased the proportion of HSCs within the ‘Interferon’ cluster from 1% to 42% (Fig 1F, p-value (DPA) < 10^-5^). The ‘Interferon’ cluster was characterized by expression of interferon response genes such as *Iigp1*, *Isg15*, *Ifit1*, *and Oasl2* (Fig 1D). *In vivo* treatment with dmPGE_2_ gave rise to a novel cluster named ‘Acute activation’ (Fig 1B) that contained 55% of dmPGE_2_-treated HSCs (Fig 1F). The cluster itself was entirely composed of dmPGE_2_-treated HSCs (Fig 1G) and marker genes include known cAMP-response genes such as *Fosl2* (Fig 1C and D) and the phosphodiesterases *Pde10a*, *Pde4b* and *Pde4d* (Fig 1D). G-CSF and indomethacin induced slight shifts in cell distribution compared to control (Fig 1E) but did not significantly alter cell proportions between clusters (Fig 1F, p-value (DPA) > 0.05 for all clusters). In conclusion, at baseline HSCs were equally distributed between three main transcriptional states, here defined as the ‘Quiescent’, ‘Activated’ and ‘Metabolism’ (Fig 1F) with few HSCs residing in ‘Interferon’ and ‘Cell cycle’ states. A two-hour *in vivo* pulse with poly(I:C) or dmPGE_2_ significantly altered distributions of HSCs between pre-existing transcriptional states and, in the case of dmPGE_2_, allowed for a novel transcriptional state to surface.

**Figure 3:**
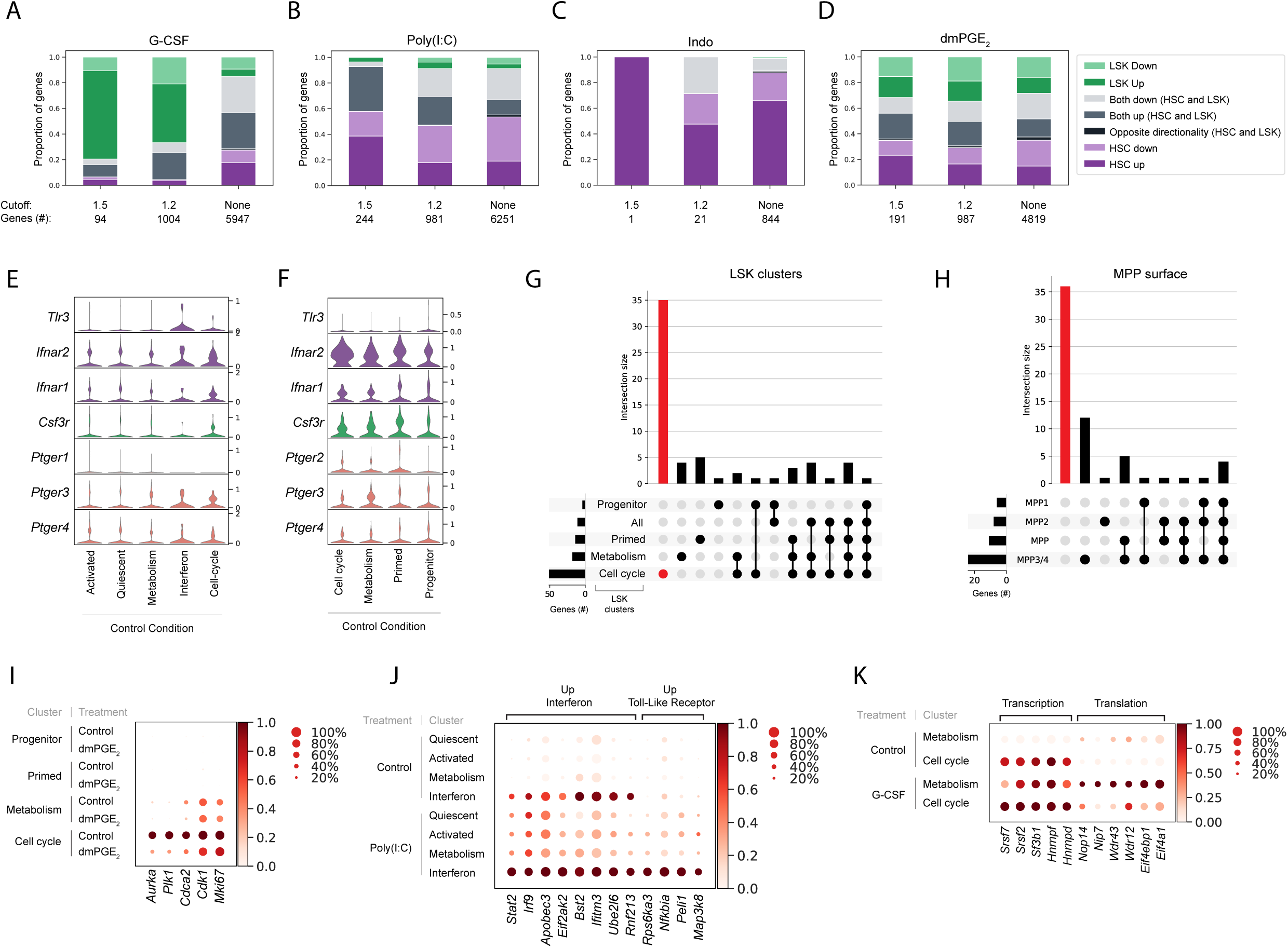
LSK and HSC cluster-specific differential gene expression analysis reveals novel, specific effects on HSC and LSK subpopulations. (A-D) Stacked bar graphs with proportion of differentially expressed genes that are unique for HSCs (purple) MPPs (green) or overlapping (grey) for G-CSF (A), Poly(I:C) (B), Indo (C) and dmPGE_2_ (D) treatment. Below each bar graph the total number of differentially expressed genes (‘Genes #’) for each fold-change (‘Cutoff’) is listed. (E-F) Violinplots of receptor expression in control HSCs (E) and LSKs (F) split by cluster. (G) Upsetplot visualizing differentially expressed genes between dmPGE_2_ and control for each cluster (horizontal bars) and intersection of gene sets between clusters (vertical bars) indicated by dots. Red bar and dot indicate specific genes in the Cell cycle cluster. (H) Upsetplot visualizing differentially expressed genes between dmPGE_2_ and control for each MPP surface phenotype. Red bar indicates genes that are absent from any of the conditions (MPP, MPP1, MPP 2, MPP3/4). (I) Dot plot of representative cell cycle genes and expression in LSK clusters in control and dmPGE_2_. (J) Dot plot of representative genes from Poly(I:C) treated and control HSC clusters (scaled expression across columns). (K) Dot plot of representative genes from the G-CSF treated and control HSC clusters (scaled expression across columns).

**Figure 4:**
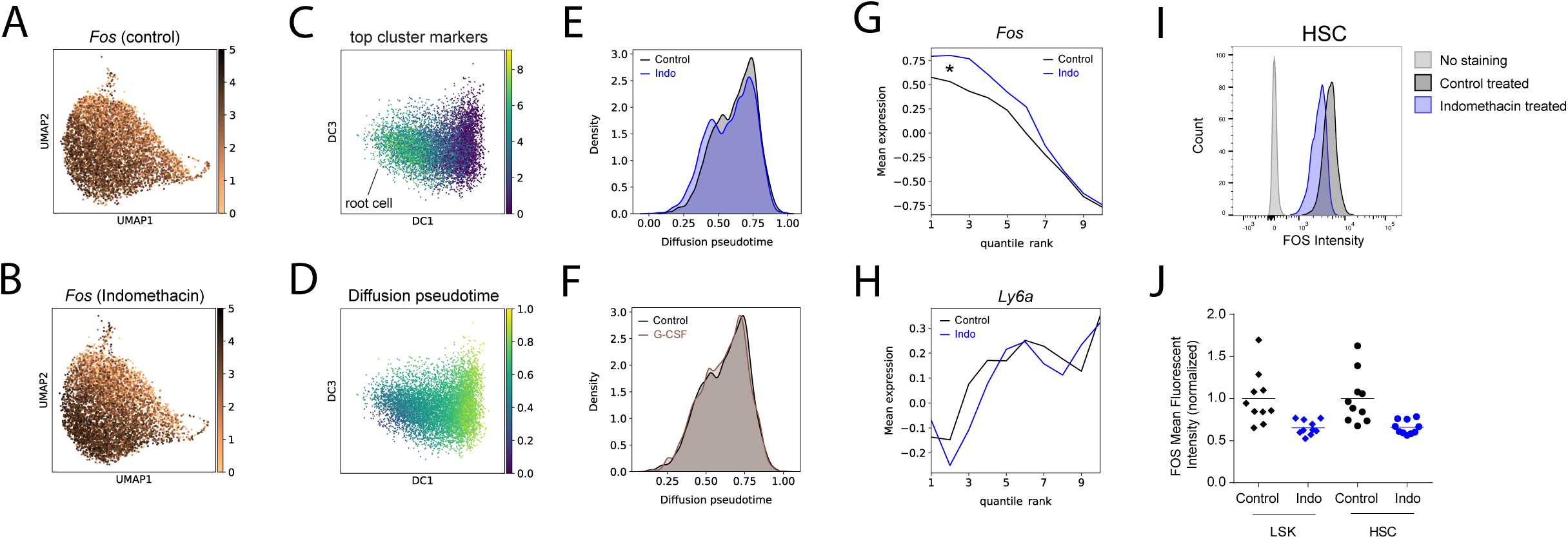
Indomethacin treatment induces change in IEG transcriptional state in HSCs. (A-H) Diffusion pseudotime analysis. UMAP plot of *Fos* expression in control (A) and upon indomethacin (B) treatment. Diffusion map embedding with combined expression of activated genes to select root cell (C) and cells colored by pseudotime (D). Kernel density of cell pseudotime distribution comparing indomethacin and control (E) and G-CSF and control (F). Average expression of *Fos* (G) and *Ly6a* (H) across cells ranked by pseudotime (cells split into 10 bins to decrease noise), change in transcript levels indicated by asterisk in G. (I) Histogram of FOS levels via intracellular FACS of HSCs, ‘no stain’ is FACS negative control, ‘control’ is FOS in untreated mice. (J) Normalized mean fluorescence intensity (MFI) for FOS in control and indomethacin treated HSCs (p = 6.2 * 10^-3^, Welch corrected t-test) and LSK cells (p = 6.6 * 10^-3^, Welch corrected t-test) across two independent biological replicate experiments. n (mice) = 20.

### Evaluation of scRNA-Seq phenotypes in entire LSK compartment to identify HSC specific cell states

In order to evaluate specificity of transcriptional heterogeneity observed within HSCs and their response to niche signals, we analyzed the transcriptome of the entire LSK compartment, which encompasses both HSCs and MPPs (Sup Fig 1A, 1G and Sup Fig 2A). To assess transcriptional response and cell states in phenotypically defined MPPs (Cabezas-Wallscheid et al., 2014, Oguro et al., 2013, Pietras et al., 2015) (MPP, MPP1, MPP2, MPP3/4, Sup Fig 1A) a Hashtag Oligonucleotide (HTO) labelling strategy was used that is part of the Cellular Indexing of Transcriptomes and Epitopes by Sequencing (CITE-Seq) methodology (Sup Fig 4A, Fig 2B and Methods (Stoeckius et al., 2018)). CITE-Seq enables tracking of cell surface phenotypes in scRNA-Seq data through barcoding of cells with antibody conjugated DNA-oligos. ScRNA-Seq gene expression of marker genes such as *Cd34*, *Cd48* and *Cd150 (Slamf1)* matched the surface phenotypes used for sorting of CITE-Seq barcoded MPPs, confirming that our workflow was successful (Fig 2C, Sup Fig 4B). Similar to the approach used for HSCs, we analyzed transcriptomic data from LSK cells as an aggregated set consisting of all four niche perturbations and control. We discovered a total of eight cell states, which similarly to HSCs displayed enrichment as opposed to exclusive expression of marker genes (Fig 2A). These clusters were labeled through analysis of enriched genes (Fig 2D, Sup Table 4), their composition of phenotypically defined cell populations tracked with CITE-Seq (Fig 2B and 2H) and by comparison to the earlier defined HSC clusters (Sup Fig 4C). LSK clusters most similar to the ‘Quiescent’ and ‘Activated’ HSC state were named ‘Progenitor’ and ‘Primed’, respectively. The ‘Progenitor’ cluster encompassed the majority of phenotypic HSCs and was significantly depleted of MPP3/4s compared to all other clusters (Fig 2H, DPA p-values < 0.02). Conversely, phenotypic HSCs were almost exclusively composed of ‘Progenitor’ cluster cells (Fig 2F). The location of HSCs at the edge of the UMAP plot could indicate the origin for differentiation (Fig 2B and Sup Fig 4F). LSK cells in the ‘Primed’ cluster represent a more committed state given their expression of *Cd34* and *Flt3*. Enrichment of *Cd37* and *Sox4* suggest priming towards a lymphoid fate (Fig 2D (Sun et al., 2013, Zou et al., 2018)). The two LSK clusters ‘Cell cycle’ and ‘Metabolism’ contained cells that are mitotically active or are on the verge of entering the cell cycle, respectively. Comparison of the top 100 enriched genes between clusters indicated a 33% overlap between the LSK ‘Metabolism’ cluster and the HSC ‘Metabolism’ cluster and 41% common enriched genes between the LSK ‘Metabolism’ cluster and the HSC ‘Cell-cycle’ cluster (Sup Fig 4B - C and Fig 2D). The myeloid cluster is defined by expression of genes such as *Mpo, Ctsg*, *Fcer1g* and *Cebp*α (Fig 2D). Consistent with previous reports (Pietras et al., 2015), MPP2s contained the highest proportion (7.6%) of myeloid cells and comprise 28% of the entire ‘Myeloid’ cell cluster (Fig 2F and 2H). Control-treated LSKs were distributed amongst four main clusters, those being ‘Primed’, ‘Progenitor’, ‘Metabolism’ and ‘Cell cycle’, that together encompassed 99% of control LSK cells (Fig 2E). Similar to our observations in HSCs, only G-CSF and indomethacin treatment did not alter cellular distributions between LSK clusters (Fig 2E, Sup Fig 4G). Conversely, dmPGE_2_ and poly(I:C) gave rise to novel clusters that were absent in control LSKs (Fig 2E and G). These treatment-induced states displayed transcriptional profiles that were similar to the HSC equivalents (Sup Fig 4B and C, Fig 2D). Surprisingly, and in contrast to HSCs, no ‘Interferon’ responsive cell state was present in LSKs at baseline (Fig 2E). Poly(I:C) treatment induced two interferon responsive clusters in LSKs, of which one showed higher mitotic activity (‘Interferon cell cycle’, Fig 2D and Sup Fig 4C). In summary, single-cell transcriptome analysis of a CITE-Seq validated LSK compartment revealed an increased proportion of lineage-committed and mitotically active and an absence of steady-state interferon responsive cells compared to HSCs. The changes in cell distribution between clusters upon niche stimulation were highly similar between LSKs and HSCs. Therefore using this analytic approach we did not uncover any HSC-specific cellular behaviors upon signaling from the niche.

### Within cluster analysis detects subpopulation specific effects of niche perturbations

To assess niche-induced transcriptional changes in greater detail we used “Model-based Analysis of Single-cell Transcriptomics” or MAST (see Methods, (Finak et al., 2015)). This approach allowed us to perform differential expression analysis on stimulants compared to the control within each cell state cluster. We compiled differentially expressed genes (DEGs) of the four stimulant conditions at three levels of expression changes across all clusters, that is using a 1.5-fold change, 1.2-fold change, and no fold-change cutoff (FDR < 0.01 see Methods and Sup Table 5). We then aggregated genes based on common (‘up/down overlap’) or unique expression (‘up/down HSC/LSK only’) within HSCs or LSKs (Fig 3A-D). G-CSF treatment perturbed gene expression more strongly within LSKs (green bars, Fig 3A) whereas interferon stimulation by Poly(I:C) predominantly affected HSCs (purple bars, Fig 3B). Receptor expression did not completely explain this difference since both the G-CSF receptor *Csf3r* and the type I interferon receptors *Ifnar1* and *Ifnar2* were expressed in a higher proportion of LSK cells compared to HSCs (Fig 3E and F). Indomethacin was found to selectively affect HSCs but overall very few genes were differentially expressed (Fig 3C). dmPGE_2_ led to a balanced effect on HSCs and LSKs, with neither compartment dominating the DEGs (Fig 3D). We further analyzed DEGs that were unique for specific clusters within either HSCs or LSKs. dmPGE_2_ stimulation decreased expression of genes that promote the cell cycle, such as *Aurka*, *Plk1* and *Ki67*, within the LSK ‘Cell cycle’ cluster (Fig 3I, 3G; in red, Sup Fig 6C). As a comparison and to mimic a dataset that could be obtained by bulk RNA-Seq after sorting of MPPs, we analyzed DEGs in MPP surface phenotypes that were not split into distinct transcriptional clusters. DEG analysis within MPP surface phenotypes failed to recover the dmPGE_2_-mediated regulation of cell cycle genes (Fig 3H, in red). The differentially expressed cell cycle genes were likely not be detected in the pseudo-bulk MPP populations since only 6.4% (MPP) to 22% (MPP2, Fig 2F) of cells belong to the ‘Cell cycle’ cluster. A similar ‘dilution’ of treatment-specific effects within the ‘Cell cycle’ cluster also occurred following G-CSF treatment when comparing pseudo-bulk and cluster separated LSK cells (Sup Fig 4D and E, in red). These results illustrated that within-cluster-based differential expression analysis is highly sensitive in identifying genes specific to particular transcriptional states within a cellular compartment. In contrast to between-cluster cell distribution analysis, the within-cluster approach revealed specific transcriptional responses to niche signals in HSCs and LSKs.

### Endogenous cell states distinguish TLR- and IFN-specific responses of Poly(I:C) treatment

To identify distinct patterns of regulation for different stimulations and hematopoietic stages, we selected genes that were differentially expressed within at least one HSC or LSK cluster. We then averaged expression of all cells within a cluster and grouped single cell clusters and genes by hierarchical clustering for HSCs (Sup Fig 5, Sup Table 6) and LSKs (Sup Fig 6, Sup Table 7). Poly(I:C) binds to Toll-like receptor 3 (TLR3) (Alexopoulou et al., 2001) which leads to expression of Type 1 interferons (IFNα and IFNβ) that in turn signal via IFNα/β receptor 1 (*Ifnar1*) and 2 (*Ifnar2*) heterodimers, all of which are expressed in HSCs (Fig 3E). We identified two expression patterns in poly(I:C)-treated HSCs that are consistent with toll-like receptor and interferon receptor signaling. The first expression pattern is driven by induction of poly(I:C) responsive genes across all cell states. In addition, these genes were also specifically enriched in the ‘Interferon’ cluster already prior to poly(I:C) stimulation (Fig 3J; ‘Up Interferon’). Genes within this group are either directly downstream of type 1 interferon receptors, such as *Stat2* and *Irf9,* or act as effector proteins involved in viral interferon response such as *Apobec3* and *Eif2ak2* (Fig 3J, Sup Fig 4A). The high expression of several interferon-induced viral response genes (*e.g. Bst2*, *Ifitm3, Ube2l6,* and *Rnf213*) in the control ‘Interferon’ cluster might point to a state of general surveillance for viral infection at baseline (Fig 3J, Sup Fig 5A). The second expression pattern (‘Up Toll-Like Receptor’) results from genes that are induced by Poly(I:C) treatment, especially in the poly(I:C) ‘Interferon’ cluster, but show low expression at baseline, even in the control ‘Interferon’ cluster. (Fig 3J, Sup Fig 5A). Genes within this signature include *Nfkbia*, *Peli1*, *Map3k8*, and *Rps6ka3* and are part of TNFα and Toll-like signaling pathways. This expression profile might therefore represent a more direct response of poly(I:C) interaction with Tlr3. Comparison of differential expression patterns across cell states allowed us to distinguish between poly(I:C)-mediated TLR- and interferon-based signaling.

**Figure 5:**
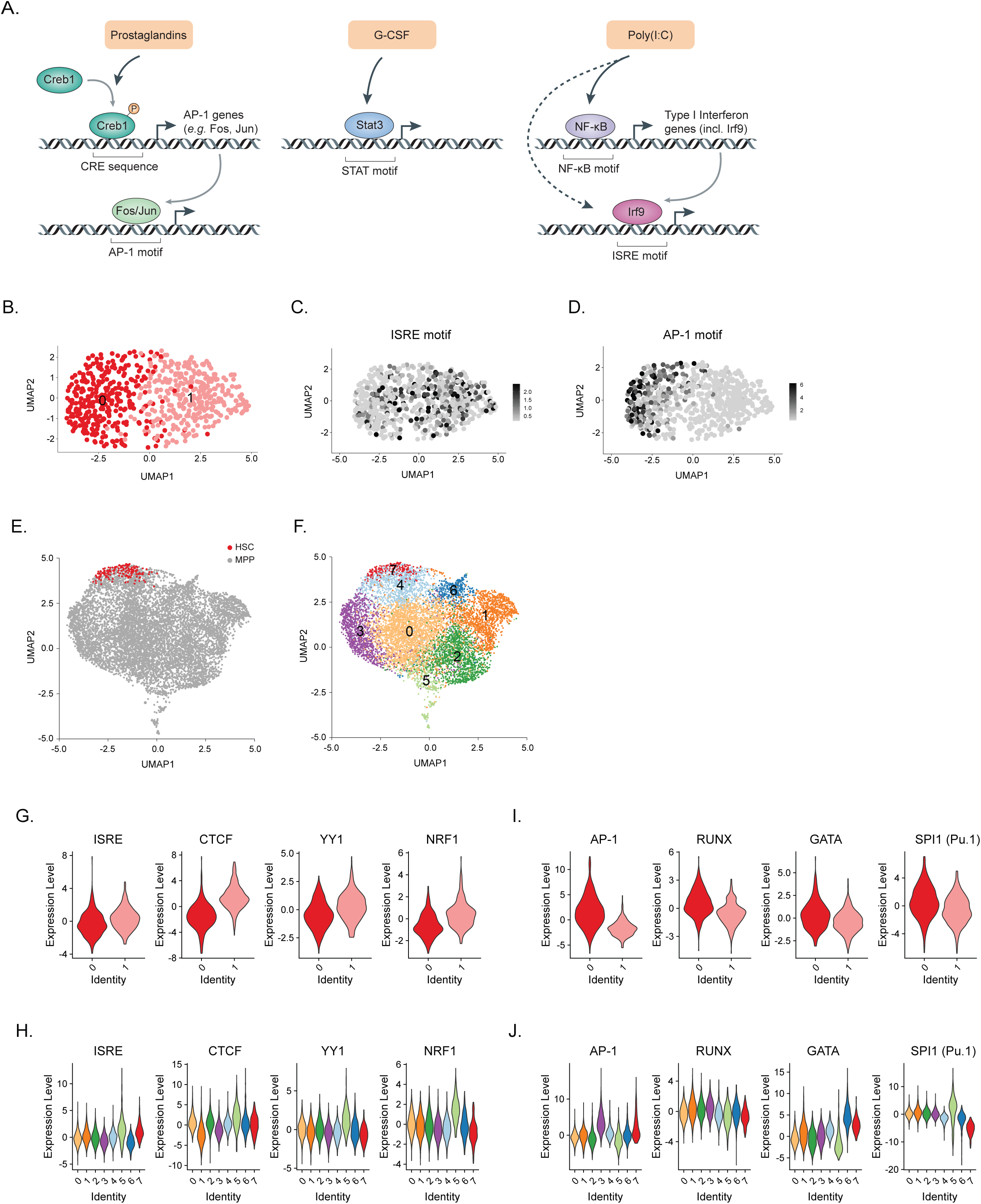
Specific transcription factor binding motif cooccurrences in HSCs. (A) Schematic of downstream transcriptional signaling pathways for niche stimulants. (B-D) UMAP plot of HSC scATAC-Seq clusters (n = 730 cells) (B) and ISRE (C) or AP-1 (D) TF motif scores. (E-F) UMAP plot of LSKs (n = 10750 cells) colored by surface phenotype (E) and scATAC-Seq clusters (F). (G-H) Violin plots of TF motif scores enriched in HSC cluster 1 in HSCs (G) and LSKs (H). (I-J) Violin plots of TF motif scores enriched in HSC cluster 0 in HSCs (I) and LSKs (J).

### G-CSF triggers changes within a metabolic transcriptional state without changing cell distribution between HSC and LSK clusters

G-CSF has been identified as a potent enhancer of granulocyte and neutrophil differentiation (Metcalf and Nicola, 1983). It is widely used as a mobilizing agent for hematopoietic stem cells in a therapeutic setting (Bendall and Bradstock, 2014). In line with its clinical use, we found niche adhesion receptors such as *ckit* and *Cd9* (Leung et al., 2011) to be downregulated following *in vivo* G-CSF treatment (Sup Fig 5B, purple arrows).

Overall G-CSF induced most DEGs within the HSC ‘Metabolism’ cluster. Hierarchical clustering revealed that G-CSF treatment alters the expression profile of the HSC ‘Metabolism’ cluster in a way that makes it more similar to ‘Cell cycle’ states (Sup Fig 5B). This resemblance is driven by induction of genes related to transcription, such as RNA binding proteins (*Hnrnpd, Hnrnpf, Hnrnpa2b1*), as well as splicing factors (*Srsf7, Sf3b1, Srsf2*) (‘Transcription’, Fig 3K). G-CSF also increased expression of transcripts involved in translation, including genes involved in ribosome biogenesis (*Nop14, Nip7, Wdr43, Wdr12*) and translation initiation (*Eif4a1, Eif4ebp1*). These transcripts were not expressed in the ‘Cell cycle’ state at baseline (‘Translation’, Fig 3K) and might be related to G-CSF-induced fate commitment towards differentiation. Overall, a two-hour pulse of G-CSF pushed HSCs towards a more metabolically active state. Our scRNA-Seq data are consistent with the original description of G-CSF as a growth factor that regulates myeloid differentiation and indicates early transcriptional responses leading to HSC mobilization.

### Endogenous Prostaglandins regulate immediate early genes within ‘Activated’ cell states

To investigate signaling from the niche in a more physiological setting, we orally treated mice for one week with indomethacin to deplete endogenous prostaglandins. Differential expression analysis identified only 21 genes (1.2-fold change cutoff) to be differentially expressed compared to the control (Fig 3A). Of the upregulated genes, 10 out of 12 can be classified as IEGs (e.g. *Fos, Fosb, Jun, Klf4 or Klf6*) (Sup Fig 5D). While cell proportions did not change between the HSC clusters (Fig 1F), distribution of cells shifted slightly towards the periphery within the UMAP plot (Fig 1E). This change in distribution was seen with individual cluster marker genes such as *Fos* and other IEGs (Fig 4A, 4B and Sup Fig7A, 7 B). To further investigate the influence of endogenous prostaglandin depletion on cell distribution while taking the entire transcriptional landscape into account, we computed diffusion pseudotime (DPT) (Haghverdi et al., 2016) between the ‘Activated’ and ‘Quiescent’ cluster in HSCs. The cell with the combined highest expression of the three top cluster markers for the ‘Activated’ state (Fig 4C, see Methods) was set as root cell and DPT was calculated originating from that root cell (Fig 4D). Indomethacin-treated cells displayed a significant shift to the left in overall pseudotime kernel density distribution (KDE), which is indicative of overall lower pseudotime (Fig 4E, p-value = 5.8*10^-12^ by Mann–Whitney U-test). No shift was observed when comparing the control to G-CSF treated HSCs (Fig 4F, p-value = 0.18). Ranking cells for each treatment condition according to pseudotime and averaging gene expression in 10 equally sized bins (quantile rank 1-10) further illustrated the change in expression of *Fos* and other IEG genes following indomethacin, especially at lower pseudotimes (Fig 4G and Sup Fig 7C; indicated by asterisks). Genes that were not part of the activated gene signature, such as *Ly6a*, did not follow the same pattern (Fig 4H), nor was a similar trend observed in response to G-CSF treatment (Sup Fig 7D). The pseudotime analysis of the scRNA-Seq data indicated a specific shift in IEG transcriptional state upon depletion of endogenous prostaglandins. To further confirm the effect of endogenous prostaglandins on IEGs in an orthogonal assay, we measured single cell protein levels by intracellular flow cytometry for FOS. Across two independent experiments, a seven day *in vivo* indomethacin treatment led on average to a 34% (SD = 8.2%) reduction in FOS mean fluorescent intensity (MFI) in HSCs (p-value = 6.2 * 10^-3^, t-test with Welch’s correction) and a mean 35% (SD = 8.6%) decrease in LSKs (p = 6.6 * 10^-3^, Fig4I and J). Overall, endogenous prostaglandin levels impact both the transcriptional state and protein levels of FOS and potentially other IEGs.

### HSC-specific chromatin architecture is an intrinsic regulator of differential response to niche signals

To better understand HSC intrinsic factors regulating the transcriptional ‘receptiveness’ to signals, we assessed chromatin states using single cell ATAC-Seq (scATAC-Seq, see Methods) of sorted HSCs and MPPs. We clustered cells based on chromatin accessibility in HSCs (Fig 5B) and LSK cells consisting of MPPs and HSCs (Fig 5E and F, Methods). To gain insight into the nature of the differentially accessible chromatin regions, we computed a per-cell transcription factor (TF) motif activity score using ChromVar (Schep et al., 2017) and evaluated enrichment of these scores across clusters. The motif activities of transcription factors STAT3, NF-κB, and CREB1 that are immediately downstream of G-CSF, Poly(I:C), and Prostaglandins (Fig 5A), respectively, were homogeneously distributed in both HSCs and LSKs (Sup Fig 8A, 8B, and Sup table 8). This result suggested that all cells have an equally responsive potential to these niche signals based on their accessible chromatin states. We did detect differential enrichment of motifs for transcription factors that are further downstream in the response to niche signals. Specifically, we found differential enrichment of the interferon signaling response element (ISRE) motif activity score in cluster 1 (Fig 5C) and AP-1 motif score in cluster 0 (Fig 5D). Interferon-regulatory factors (IRFs) that bind ISREs are induced by NF-κB signaling as well as direct targets of Poly(I:C) intracellular binding ((Negishi et al., 2018), Fig 5A). In addition to IRFs, HSC cluster 1 is characterized by motifs for transcriptional regulators that are important for cellular metabolism, cell growth, and differentiation such as CTCF, YY1, and NRF1 (Fig 5G and Sup Fig 8C). Enrichment of a similar set of motifs in LSK cell cluster 5 (Fig 5F and H in light green, Sup Fig 8G) indicated that this motif combination is not unique for HSCs. The location of cluster 5, being most distally from phenotypic HSCs on the UMAP plot (Fig 5E), further suggested that cells in this chromatin state are committing to or undergoing differentiation. The AP-1 motif which is specifically enriched in HSC cluster 0 can be bound by FOS and JUN, both downstream effectors of the Prostaglandin/CREB1 signaling pathway ((Luan et al., 2015), Fig 5A). We furthermore found motif activity enrichment in HSC cluster 0 for several key HSC lineage-specific master transcription factors including RUNX, GATA, and PU.1 (SPI1) (Fig 5I and Sup Fig 8D) as well as SMAD, another signal-responsive transcription factor (Sup Fig 8E and F). In contrast to IRFs, LSK cells did not contain a corresponding cluster where the same motifs cooccur as in HSC cluster 0 (Fig 5J and Sup Fig 8H). Therefore, the chromatin features of cluster 0 may be specific for HSCs. We found no indication that differential responsiveness to the niche stimulants is a result of distinct chromatin states of transcription factor motifs directly downstream of these signals. Rather, our analysis implicated cell intrinsic heterogeneity of further downstream effectors, such as AP-1 and IRFs. The specific motif co-occurrence of AP-1 and HSC-lineage specific master factors, combined with their absence in LSKs, indicates an HSC-specific chromatin state.

## Discussion

Here, we provide the first transcriptional and epigenetic single cell analysis of a highly purified, functionally validated HSC population. Our work reveals that highly purified HSCs exist in fluent transcriptional and epigenetic states rather than distinctly separated cell types. Niche perturbations rapidly shift HSC distribution between HSC states within hours of signaling, providing evidence that the transcriptional states are highly dynamic and allow HSCs to quickly transition between states in response to different stimuli. Our single cell chromatin studies indicate cell intrinsic HSC heterogeneity that predisposes subpopulations for certain transcriptional responses. We detected an HSC specific co-occurrence of signaling and lineage-specific transcription factor motif activities that is consistent with our previous observation in human hematopoietic progenitors (Trompouki et al., 2011, Choudhuri et al., 2020)). Absence of similar chromatin features in LSK progenitors may implicate a link to some of the unique functional capacities of HSCs, such as self-renewal. Overall, our data indicates that the single cell landscape of *in vivo* derived, functional HSCs is made up of a unique chromatin architecture with fluent transcriptional states, some of which can be rapidly influenced by signals from the niche.

Our combined scRNA-Seq and CITE-Seq approach allowed us to gain insights into the transcriptional landscape of HSCs and phenotypically defined MPP populations within the LSK compartment at steady state and following niche perturbations. Our results enabled us to connect the transcriptional profile on a single cell level to the previously described phenotypic behaviors of these MPP populations (Oguro et al., 2013, Pietras et al., 2015, Cabezas-Wallscheid et al., 2014). In addition, the CITE-Seq approach allowed us to cross-validate our transcriptional cell state assignments. The HTO CITE-Seq method provided a flexible tool to evaluate and compare transcriptional profiles within phenotypically defined populations because the technology used here is not dependent on the availability of specifically conjugated antibodies against particular surface receptors. In addition, *Xist* expression was used to deconvolute pooled male and female cells. While our analysis revealed only a weak sexual dimorphism that is consistent with previous reports (Nakada et al., 2014, Gal-Oz et al., 2019), the negligible additional investment to obtain data from both sexes may become the default experimental design in mammalian scRNA-Seq experiments. Our work presents evidence for two value-adding pooling strategies that allow for further insights into cell populations analyzed by scRNA-Seq.

We used a two-pronged strategy to assess the specificity of niche perturbations in HSCs or LSKs. First, we determined changes of cell proportions between cell states. Second, we evaluated differential expression within particular cell states following stimulation. The strength of transcriptional perturbation could not solely be estimated based on the distribution of cells within clusters alone. G-CSF did not change the cell proportions between clusters but rather elicited strong transcriptional responses within a given cell state. Furthermore, analysis of DEGs within clusters helped tease apart interferon-versus toll-like receptor response genes induced by Poly(I:C) treatment. We also uncovered a specific effect of dmPGE_2_ that only occurs within the LSK ‘Cell cycle’ cluster. A recent study corroborates our finding that dmPGE_2_ affects cell cycle states in the bone marrow (Patterson et al., 2020). Our results show that this within-cluster differential gene expression analysis was most sensitive to reveal HSC or LSK specific responses to niche perturbations.

There is a tradeoff between the strength of a perturbation required for experimental robustness versus studying signals that are more physiologically relevant but lead to more subtle changes within and between cells. Here, we evaluated response of HSCs to three different activators mimicking niche signals that were dosed 2-4 orders of magnitude higher than what an animal would ever encounter during actual injury or infection (Eyles et al., 2008, Porter et al., 2013, Hoggatt et al., 2013, Sheehan et al., 2015). To assess niche-derived signals in a more physiological setting, we administered the Cox1/2 inhibitor indomethacin orally for one week to deplete endogenous prostaglandins. As expected, the changes in gene expression with indomethacin were much weaker than those observed after acute injection with dmPGE_2_, G-CSF, and poly(I:C). ScRNA-Seq analysis offers unique tools to evaluate gene expression changes in response to weak perturbations. Pseudotime analysis showed that depletion of endogenous prostaglandins using indomethacin leads to a small but significant shift in the transcriptional state of HSCs. The effect of indomethacin on IEGs such as *Fos* was further validated in independent FACS experiments which showed that the transcriptional programs implicated through pseudotime were also found to be perturbed using this orthogonal assay. How exactly the increase in RNA levels of *Fos* observed in scRNA-Seq can be reconciled with decreased FOS protein levels determined by FACS analysis will need to be addressed in future experiments. Another important implication and potential caveat highlighted by our findings is that RNA and protein levels may not always positively correlate, even on a single cell level. Regardless, scRNA-Seq technologies provides sensitive tools to interrogate subtle changes in cellular states.

In summary, we showed that single cell approaches provide a rich and sensitive tool to analyze transcriptional and epigenetic states of HSCs during homeostasis and upon niche perturbation. We found that HSCs exist in dynamic cell states and niche signals can induce rapid transitions between, as well as changes within, these HSC states. While our work does not reveal whether these transcriptional states are associated with specific niches *in vivo*, novel spatial transcriptomic approaches provide exciting new opportunities to address such questions (Rodriques et al., 2019). Additionally, recently developed barcoding strategies enable assessment of niche-induced transcriptional changes and functional potential of single cells within the same experiment (Rodriguez-Fraticelli et al., 2020). Understanding endogenous levels of niche-derived factors and the associated transcriptional and epigenetic responses will advance our basic understanding of stem cells and their potential applications in the clinic.

## Materials and Methods

### WET LAB METHODS

#### Mice and niche stimulant treatment

For the HSC Replicate 1 experiment we used the following mouse strain (#016617) that was obtained from Jackson labs but bred in house. For niche stimulant treatments male and female mice (8–10 weeks) were ordered from Jackson labs (strain CD 45.2 (Ly5.2), #00664). Mice were kept for at least 1 week in the animal facility before initiating experiments and allocated at random (by cage) into experimental groups. Indomethacin (Sigma, 6mg/l) was administered for 7 days in acidified drinking water to maintain stability (Curry et al., 1982, Pratico et al., 2001). Indomethacin supplemented drinking water was changed every other day. Mice were injected with the following drugs and euthanized after 2 hours: Poly(I:C) HMW (Invivogen), IP injection 10mg/kg (Pietras et al., 2014). G-CSF Recombinant Human Protein (Thermo fisher), IP injection, 0.25mg/kg (Morrison et al., 1997). DmPGE2 (Cayman), SC injection, 2mg/kg(Hoggatt et al., 2013). Mice were weighed before injection and injection volume was adjusted to ensure equal dose between individual mice. The ‘control’ condition from the niche stimulant treatments was also used as the second independent biological replicate of unperturbed HSCs (HSC Replicate 2). All animal procedures were approved by the Harvard University Institutional Animal Care and Use Committee.

#### Bone marrow preparation and Fluorescence activated cell sorting (FACS)

Whole Bone marrow was isolated from femur, tibia, hip and vertebrae via gentle crushing using a mortar and pestle. Stem and progenitor cells were enriched via lineage depletions (Miltenyi Biotech, 130-090-858). Antibodies, dilutions and vendors are listed in Sup Table 9. Cells were stained for 1.5 hours based on published best practice protocols for assessing CD34 labelling (Ema et al., 2006). HSCs (LSK, CD48-, CD150+, CD34-), MPP1s (LSK, CD48-, CD150+, CD34+), MPPs (LSK, CD48-, CD150-), MPP2s (LSK, CD48+, CD150+), and MPP3/4s (LSK, CD48+, CD150-) were sorted on a FACSAria (Becton Dickinson) and representative sorting scheme is shown in Supplemental Fig 1A. Purity of > 80% was ensured by reanalyzing each sorted population.

#### Sample size estimation and sample batching

To determine appropriate sample sizes of mice and HSCs we performed an initial experiment on fresh HSCs (HSC Replicate1) which yielded estimated number of 2382 cells (after filtering), and which resolved biologically meaningful clusters. In subsequent experiments we therefore targeted obtaining a similar or higher cell number. For niche stimulant treatment we based our sample size of 5 male and 5 female mice on this initial experiment. Because of sample processing times a maximum of two conditions could be performed on the same day, resulting in three separate days of experiments. To mitigate batch effects resulting from different experimental days the following precautions were taken. (1) All mice included in the niche stimulant treatment were ordered from the same batch from JAX. (2) Control mice were administered acidified water injected with DMSO to control for both unspecific perturbations that might result from the niche stimulant treatments. (3) All experiments were performed within less than one week and single cell libraries were prepared together for all samples after the initial droplet reaction was frozen. (4) FACS gates were set up initially but left constant for each experiment. Single color controls as well as Fluorescence minus one (FMO) controls ensured that there was minimal day-to-day technical drift on the FACS instrument.

#### Intracellular staining for FACS

BM extraction, lineage depletion and surface marker staining were performed as described above. Cells were fixed and permeabilized for intracellular staining according to manufacturer’s instructions (BD biosciences, 554714). Intracellular staining was performed for 30 minutes on ice. Samples were analyzed on an LSRII FACS analyzer.

#### Limit dilution Transplantation assay

Recipient CD45.2 (Jax #00664) mice were gamma-irradiated (Cs-137 source) with a split dose of 5.5 Gy each one day before transplantation. HSCs were isolated from CD45.1 (Jax #002014) donors and transplanted with 200 000 whole bone marrow cells (CD45.2) via retro-orbital injection. Donor cell engraftment was monitored monthly for 16 weeks using a LSRII FACS analyzer (Becton Dickinson). Flow cytometry data were analyzed with FlowJo (Tree Star). HSC frequency was calculated using the following website http://bioinf.wehi.edu.au/software/elda/.

#### Single-cell RNA and ATAC sequencing library preparation and sequencing

Male and female cells were sorted separately but pooled in equal ratios before further downstream processing. For CITE-Seq HTO labelling of MPP populations, 0.25ug of TruStain FcX™ Blocking reagent (Biolegend) was added for 10 minutes on ice. Each MPP populations was labelled with 1ug of TotalSeq™ antibody cocktail (Biolegend, see Sup Table 9) and incubated for 30 minutes on ice. After washing, cells were resuspended in small amounts, counted and pooled in equal ratios. Each drug treatment condition resulted in one pooled MPP and one HSC sample that were processed separately for scRNA-Seq according to manufacturer’s recommendations (10X Genomics, 3’ V2 for HSC Replicate1 experiment and V3 for niche stimulant treatments). Briefly, for pooled MPPs, no more than 10 000 cells were loaded. For HSCs, all sorted cells (between 2222 sorted events for dmPGE2 and 12017 sorted events for control) were loaded on the 3’ library chip. For preparation of HTO – surface libraries manufacturer’s recommendations (Biolegend) were followed. For ATAC-seq HSCs and MPPs (pooled MPP, MPP1, MPP2 and MPP3/4) were sorted as described above from 5 male and 5 female mice (strain CD 45.2 (Ly5.2), JAX strain #00664). Nuclei were isolated and libraries were prepared using manufacturers recommendations (10x Chromium Single Cell ATAC). Libraries were sequenced on a Next-seq 500, 75 cycle kit (‘Replicate 1’, scRNA-Seq) and NOVAseq 6000, 100 cycle kit (‘Replicate 2’ and niche stimulant treatments, scRNA-Seq, sc-ATAC-seq).

### COMPUTATIONAL AND STATISTICAL ANALYSES

All code and a detailed description of the analysis is available in the following Github repository https://github.com/evafast/scrnaseq_paper). To ensure reproducibility the entire analysis (except cellranger and Cite-seq count) was entirely performed in Docker containers. Containers used for the analysis are indicated in the Jupyter notebooks and corresponding images are available on dockerhub (repository: evafast1). Interactive cell browser web app is available here: (https://mouse-hsc.cells.ucsc.edu). Raw data are available with GEO accession code GSE165844.

#### Demultiplexing and generation of count matrices

Cellranger (v3.0.1) command ‘mkfastq’ was used to demutliplex raw base call (BCL) files into individual samples and separate mRNA FASTQ files and HTO surface fastq files. The cellranger ‘count’ command was used with default options to generate gene by cell matrices from mRNA FASTQ files. CITE-Seq count (version 1.4.3) was used to generate surface count by cell matrices from the HTO surface FASTQ libraries. For the fresh HSC Replicate1 experiment cellranger (v2.1.0) was used for demultiplexing and count matrix generation. The mm10 reference genome was used for all alignments. For scATAC-seq cellranger-atac mkfastq and count (1.2.0) was used for demultiplexing and alignment and generation of the fragment file. To generate the count matrix MACS2 was run with default parameters (keeping duplicates) on the aligned reads. Resulting peak summits were extended to 300 bp and counts were extracted from Fragment file using a custom script (see Github repository) to generate a count matrix.

#### Quality control and Filtering and dimensionality reduction of scRNA-Seq data

The main parts of the bioinformatic analysis of scRNA-Seq data was performed using the python package scanpy (Wolf et al., 2018). For filtering and quality control best practice examples were followed (Luecken and Theis, 2019). Count matrices were filtered on a gene and cell level. Cells were excluded with either less than 3,000 UMIs, less than 1,500 (LT) or 2,000 (MPPs) genes or more than 20,000 (LT) or 30,000 (MPPs) counts. A cutoff of no more than 10% UMIs aligned to mitochondrial genes per cell was applied. Genes expressed in less than 20 cells were excluded from the analysis. Counts were normalized to 10,000 per cell and log transformed. Features (genes) were scaled to unit variance and zero mean before dimensionality reduction. To reveal the structure in the data we built a neighborhood graph and used the leiden community detection algorithm (Traag et al., 2019) to identify communities or clusters of related cells (see also below). The UMAP algorithm was used to embed the high dimensional dataset in a low dimensional space (Becht et al., 2018). Differential proportion analysis (DPA) was used for comparing cell proportions between clusters as previously described (Farbehi et al., 2019). Interactive visualization app of scRNA-Seq data was prepared using UCSC Cell Browser package (Speir et al., 2020).

#### Demultiplexing of CITE-Seq hashtag data

We used the DemuxEM (Gaublomme et al., 2019) implementation in pegasuspy (https://github.com/klarman-cell-observatory/pegasus/tree/0.17.1) to assign MPP surface identities and demultiplex to the pooled MPP sample. First background probabilities (‘pg.estimate_background_probs’) were estimated using default settings and ‘pg.demultiplex’ was run adjusting the alpha and the alpha_noise parameter to maximize cell retrieval by singlet classification. Assignments were validated by plotting count matrix in UMAP space and observing four distinct clusters indicative for the four HTO labels that were pooled. The proportion of demultiplexed cells matched the original pooling ratio. Analysis of coexpression of sex specific genes allowed for further validation of the doublet rate. Proportion of cells classified by DemuxEM as doublets exceeded doublet rate estimated by co-expression of sex specific genes.

#### Batch correction

Because of timing required for sort and sample prep it was impossible to sort HSCs from all conditions on one day (see also ‘Sample size estimation and sample batching’ above). To correct for potential batch effects we used ComBat (Johnson et al., 2007) with default settings on the log2 expression matrix, allowing cells to be clustered by cell type or cell state. Batch correction results were similar when we used Scanorama (Hie et al., 2019). To correct for potential sex specific differences Xist counts were regressed out. Raw data was used for all differential expression analysis and plotting of single cell gene expression values. Batch corrected counts were used for clustering and diffusion pseudotime analysis.

#### Optimal cluster parameter selection

Since HSCs and MPPs are highly purified cell populations we did not observe any clearly separated clusters in UMAP space. To aid the optimal choice of hyperparameters for leiden clustering we used a combination of Silhouette Coefficient and Davies–Bouldin index. We first validated this approach using the PBMC3K (from 10x genomics, scanpy.datasets.pbmc3k()) silver standard dataset. We iterated through a range of KNN nearest neighbors and Leiden resolution combinations measuring average Silhouette coefficient and Davies-Bouldin index in PCA space for each combination. Plotting the optimal value for Silhouette score and Davies-Bouldin index versus increasing numbers of clusters allowed for the determination of appropriate cluster number for the dataset. For the PBMC dataset there was a clear drop-off in optimal value after 8 clusters, which is corroborated by most single cell tutorials that also report 8 clusters for this dataset. After validation of this approach on PBMCs we used assessed Silhouette Coefficient and Davies–Bouldin index for different clustering results of our own HSC and MPP datasets. This allowed us to select the optimal hyper-parameters for each cluster number. The approach was validated by using data driven parameters to compare two independent biological replicates of control HSCs (‘Repl_1’ and ‘Repl_2’).

#### Differential expression using MAST

Differential expression analysis was performed using MAST (“Model-based Analysis of Singlecell Transcriptomics, (Finak et al., 2015). This method is based on a Hurdle model that takes into account both the proportion of cells expressing a given transcript as well as transcript levels themselves while being able to control for covariates. Based on previous reports differential expression cutoff was set at 1.2 fold (Smillie et al., 2019) and a more stringent cutoff of 1.5 fold was also included. Only genes that were expressed in at least 5% of the cells were considered for differential expression analysis. FDR (Benjamini & Hochberg) cutoff was set at 1%. For drug treatments differential expression between treatment and control was assessed within the entire MPP or LT dataset and within each cluster controlling for number of genes per cell and sex. For differential expression analysis between male and female cells at baseline, control datasets were analyzed with clusters and number of genes as a co-variates. For sex specific effects of drug treatments samples were split by sex and analyzed separately. Resulting differential expression coefficients were compared between male and female cells. To identify gene signatures with common patterns, for each treatment average expression of differentially expressed genes was extracted per cluster, scaled (z-score) and grouped together by similarity using hierarchical clustering (seaborn.clustermap, Euclidian distance, single linkage).

#### Diffusion pseudotime analysis

For diffusion pseudotime analysis (Haghverdi et al., 2016) cells from the ‘Quiescent’ and ‘Activated’ cluster were selected for the following treatments: control, indomethacin and G-CSF. We recalculated PCA and UMAP embeddings in this reduced dataset. Re-clustering using the Leiden algorithm was used to exclude outlier cells and assess top enriched genes within the new ‘Activated’ cluster. Raw expression of the three top enriched genes (Nr4a1, Nr4a2, Hes1) was summed to robustly select the most highly ‘activated’ cell as a root cell. Diffusion pseudotime was calculated with the following function in scanpy (‘sc.tl.dpt’) using default settings. Cells were ranked according to pseudotime and kernel density distribution was plotted using a bandwidth of 0.02. The Mann-Whitney U test was used to assess if cells from different samples are drawn from the same pseudotime distribution. To analyze gene expression across pseudotime, for each sample cells were split into ten equally sized bins according to ascending pseudotime. Bin 1 contained the first 10% of cells with the lowest pseudotime and bin 10 contained the 10% of cells with the highest pseudotime. Average gene expression for representative genes were plotted for each bin and sample.

#### Single cell ATAC-seq

The R package Signac (https://github.com/timoast/signac<u>, version 0.2.5</u>), an extension of Seurat (Stuart et al., 2019), was used for quality control, filtering of ATAC-seq peaks counts and plotting. Quality of scATAC dataset was ensured by presence of nucleosomal banding pattern and enrichment of reads around transcription start sites (TSSs). Cells were removed with a less than 1 000 or more than 20 000 fragments in peaks. Male and female cells were classified according to absence or presence of Y-chromosome reads. Since distribution of male and female cells appeared uniform across all analyses no downstream correction was taken for sex. Term frequency-inverse document frequency (TF-IDF) was used for normalization and dimensionality reduction was performed by singular value decomposition (SVD). Cells were clustered using the Louvain community finding algorithm after a neighborhood graph was built with k = 20 (HSCs) or k = 30 (LSK) nearest neighbors. To calculate TF motif scores, ChromVAR (Schep et al., 2017) was run with default parameters using the JASPAR 2018 motif database.

## Supplemental Tables

**Supplemental Table 1: Sequencing metrics**

Sequencing metric output from cellranger.

**Supplemental Table 2: Overlap of differentially regulated genes in male and female HSCs and LSKs**

Table summarizing number of differentially regulated genes within male and female HSCs and LSKs. Over-representation analysis odds ratio and p-value were calculated using a Fisher’s exact test.

**Supplemental Table 3: Differential expression result (MAST) by sex**

Each tab contains a treatment vs. control comparison (dmPGE_2_, Indo, Poly(I:C), G-CSF). Each cluster was compared to its respective control cluster separated by sex. Log fold change and adjusted p-value from the Hurdle model are listed for genes with p-values < 0.01.

**Supplemental Table 4: Marker gene enrichments in scRNA-Seq clusters**

Marker gene enrichment was calculated using a Wilcoxon-Rank-Sum test. Score (suffix ‘_s’) indicates the z-score of each gene on which p-value computation is based. Other fields are logfold change = suffix ‘_l’ and False discovery adjusted p-value – suffix ‘_p’.

**Supplemental Table 5: Differential gene expression result (MAST)**

Each tab contains a treatment vs. control comparison (dmPGE_2_, Indo, Poly(I:C), G-CSF). Each cluster was compared to its respective control cluster. Log fold change and adjusted p-value from the Hurdle model are listed for genes with p-values < 0.01.

**Supplemental Table 6: Average expression per cluster of differentially regulated genes in HSCs**

Count normalized and log transformed UMI counts from were averaged across HSC clusters for differentially regulated genes from MAST

**Supplemental Table 7: Average expression per cluster of differentially regulated genes in LSKs**

Count normalized and log transformed UMI counts from were averaged across LSK clusters differentially for regulated genes from MAST.

**Supplemental Table 8: ChromVar TF motif activity score enrichment in LSK and HSC scATAC clusters**

ChromVar motif activity score enrichment for HSC and LSK scATAC clusters.

**Supplemental Table 9: List of antibodies used**

## Acknowledgements

The authors thank members of the Zon, Wagers, Camargo and Scadden lab for helpful technical and scientific discussions, Serine Avagyan and Elliott Hagedorn for critical reading of the manuscript, Sai Ma for scATAC-Seq analysis advice and critical reading of the manuscript, Maximilian Haeussler and Matthew Speir for help with setting up and hosting the cell browser app and the HCBI, HSCRB FACS core, Office of Animal Resources, and the Bauer Core Facility at Harvard University for technical support. This work was supported by grants from the National Institutes of Health (P01HL131477-04, R01 HL04880, PPG-P015PO1HL32262-32, 5P30 DK49216, 5R01 DK53298, 5U01 HL10001-05, and R24 DK092760) (to L.I.Z.), the Leukemia & Lymphoma Society (Scholar grant 5372-15)(to E.M.F.) and a Boehringer Ingelheim Fonds PhD fellowship (to A.S.). L.I.Z. is an Investigator of the Howard Hughes Medical Institute.

## Authorship Contribution

E.M.F. and L.I.Z. designed the study; E.M.F., A.S. and M.M. performed the experiments and interpreted the data; E.M.F generated the figures and A.S. edited the figures; N.B. and J.G. provided technical guidance for single cell RNA-Seq experiments; E.M.F., E.L., N.B. and S.Y. performed bioinformatic analysis; E.M.F and Y.Z. supervised the bioinformatic analysis; F.C. and D.T.S. provided insights on the analysis and interpretation of the data; E.M.F. wrote the manuscript; A.S., M.M. and L.I.Z. revised the manuscript; all authors edited the manuscript; and L.I.Z. provided funding support.

## Supplemental Figure legends

**Supplemental Figure 1:**
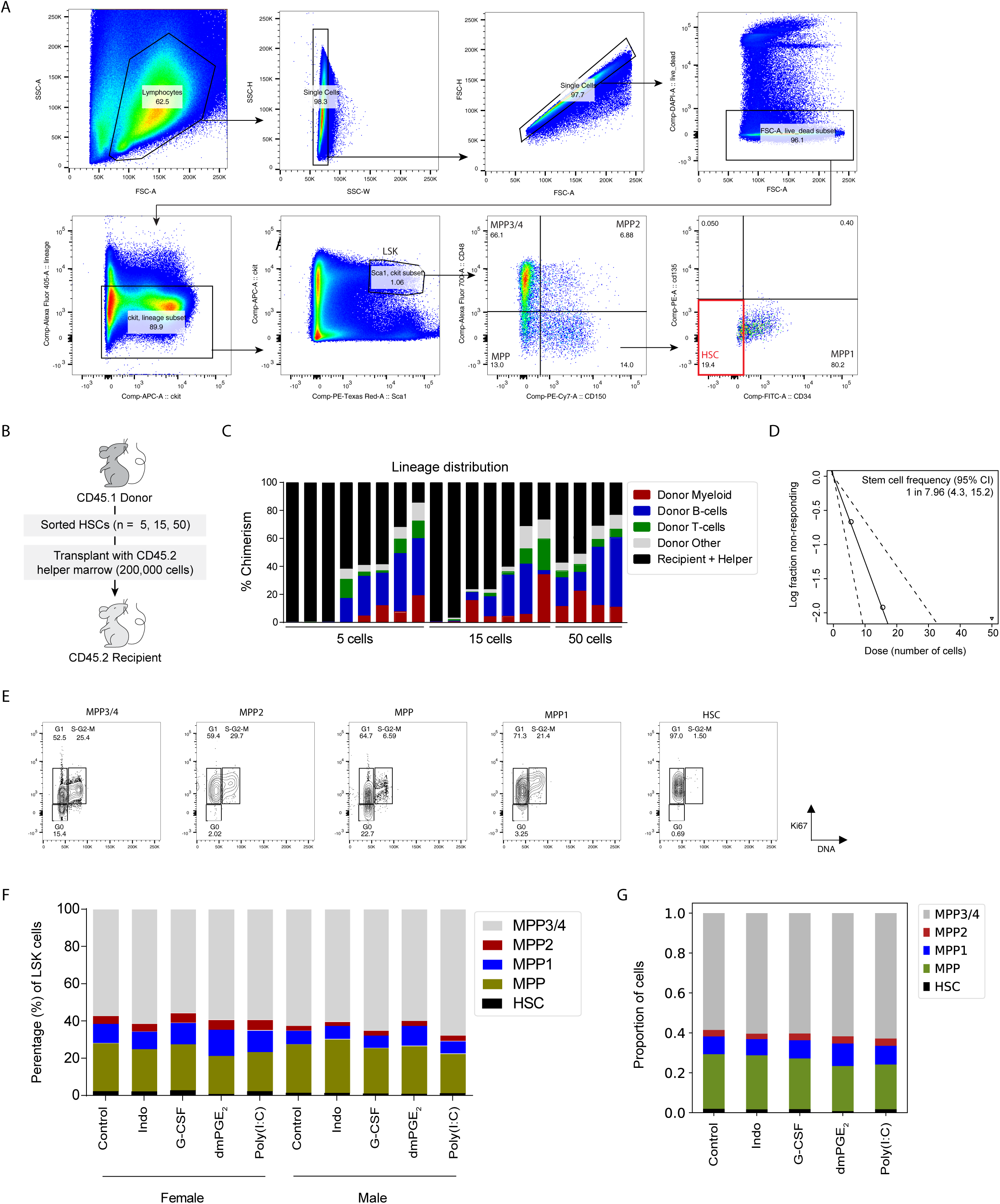
Functional characterization of HSC populations confirms high regenerative capacity. (A) Sorting scheme of MPPs and HSCs. Cells were lineage depleted prior to sort. (B) Schematic of limit dilution transplant experiment. (C) Chimerism per mouse and lineage distribution 4 months post-transplant. (D) Limit dilution analysis for transplant experiment. (E) Cell cycle FACS in MPP and HSC populations. (F) Percentage of MPPs and HSCs within LSK cells at baseline and after niche stimulation. (G) Proportion of each surface phenotype within all five experimental conditions after computational reassembly of the LSK compartment.

**Supplementary Figure 2:**
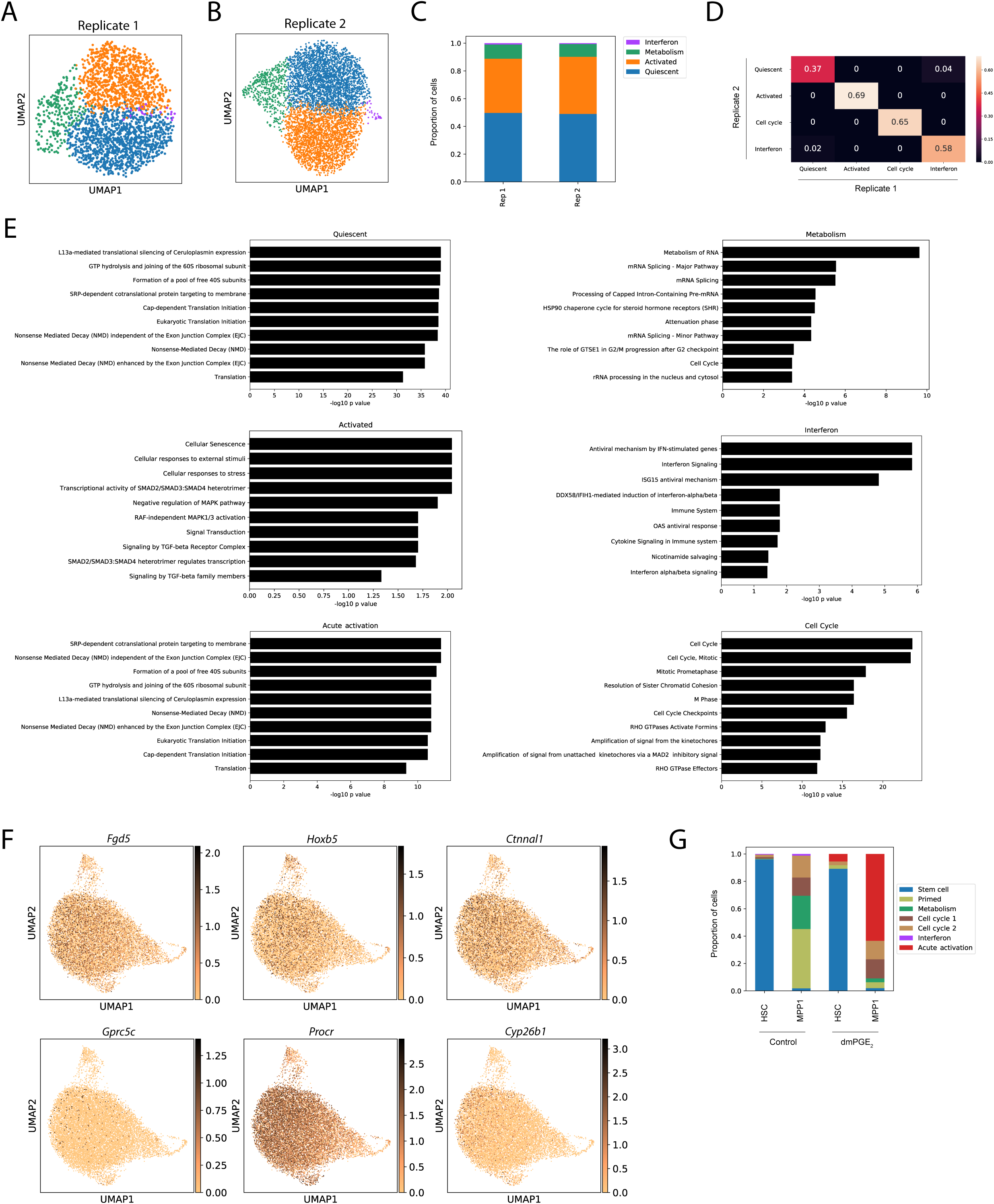
Validation of single cell RNA-Seq clustering with independent replicates, pathway enrichment and candidate genes. (A-D) Comparison of clustering of control HSCs in two biological replicates. UMAP plots for replicate 1 (A, n = 2382 cells) and 2 (B, n = 5334 cells) and summarized cell proportions in each cluster (C). (D) Pairwise comparison of the proportion of overlap of the top 100 enriched genes for each cluster. (E) Reactome pathway analysis with enriched genes for each HSC cluster. (F) UMAP plots with expression of previously described HSC markers. (G) Proportion of dmPGE2 and control cells within clusters split by surface phenotype for HSCs and MPP1s.

**Supplementary Figure 3:**
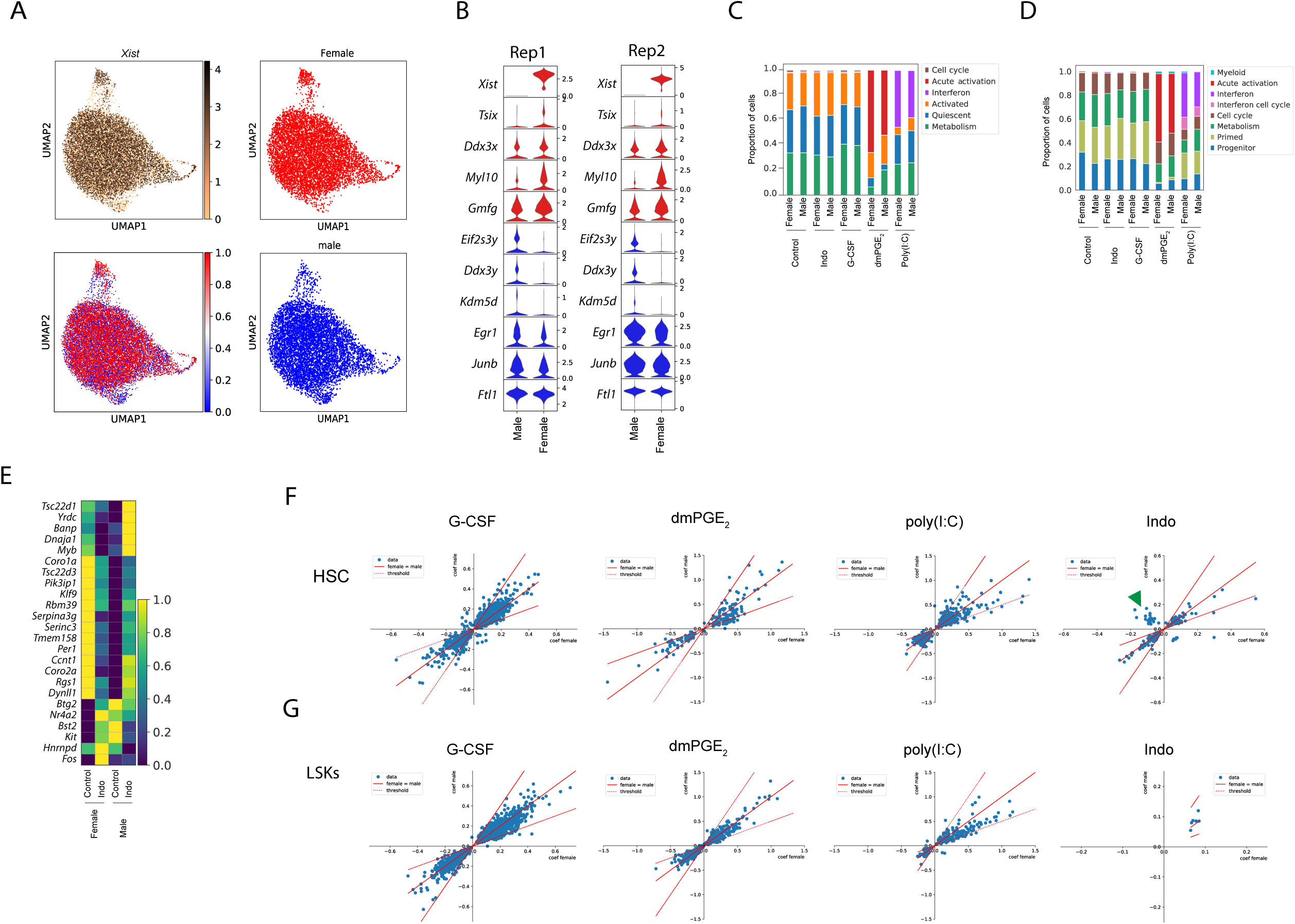
Little sexual dimorphism in HSCs in steady state and upon stimulation. (A) *Xist* expression, classification of male and female cells and male and female cells plotted separately. (B) Stacked Violin plots of all consistent sexual dimorphic genes (red; female, blue; male) for two independent biological replicates. (C-D) Proportions of male and female HSCs (C) and LSK cells (D) within clusters for each drug treatment (p(DPA) > 0.05 for all male vs. female comparisons). (E) Heat map of average expression for ‘opposite directionality’ genes in HSCs for indomethacin (‘Indo’) and control. (F) Scatter plot of differential expression coefficient (converted to log_2_ scale) induced by stimulants in HSCs (F) and LSKs (G) between male (y-axis) and female (x-axis). Solid red line indicates equal expression coefficients (coef female = coef male) and dashed line indicates a two-fold deviation (2*coef female = coef male or vice versa). Green arrowhead indicates ‘opposite directionality’ genes in indomethacin (shown also in E).

**Supplemental Figure 4:**
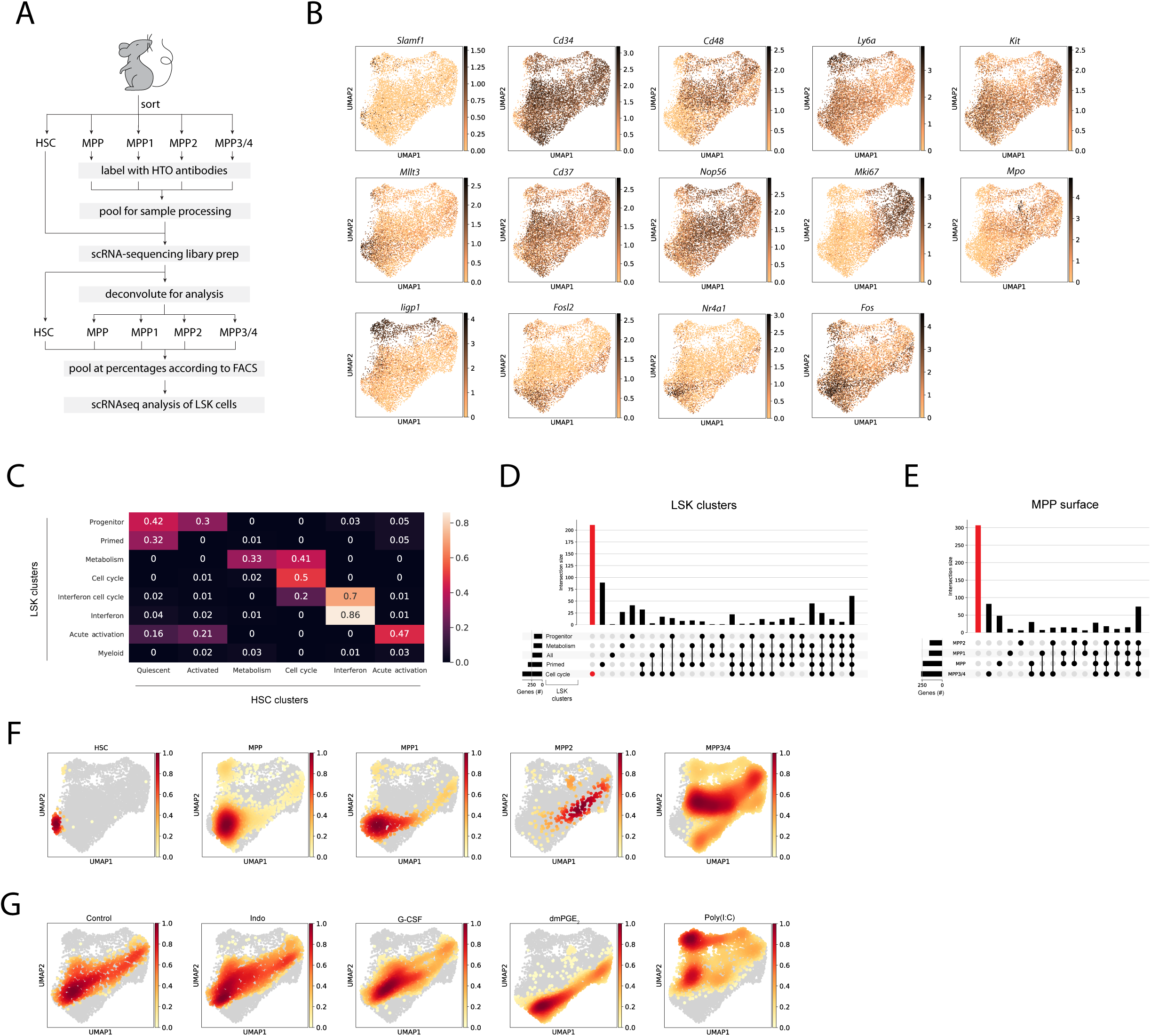
Gene expression in LSKs enables evaluation of specificity of niche stimulation in different cell populations. (A) Schematic of LSK pooling and CITE-Seq surface hashtag (HTO) methodology. (B) UMAP plots with expression of selected genes in LSK cells. (C) Heat map of proportion of overlap between the 100 top enriched genes for LSK (rows) and HSC (columns) clusters. (D) Upsetplot visualizing differentially expressed genes between G-CSF and control for each cluster (horizontal bars) and intersection of gene sets (vertical bars) between clusters indicated by dots. Red bar and dot indicate specific genes in the Cell cycle cluster. (E) Upsetplot visualizing differentially expressed genes between G-CSF and control for each MPP surface phenotype. Red bar indicates genes that are absent from any of the conditions (MPP, MPP1, MPP 2, MPP3/4). (F-G) UMAP density graphs visualizing the distribution of cells by surface phenotype (F) or by niche stimulation (G).

**Supplemental Figure 5:**
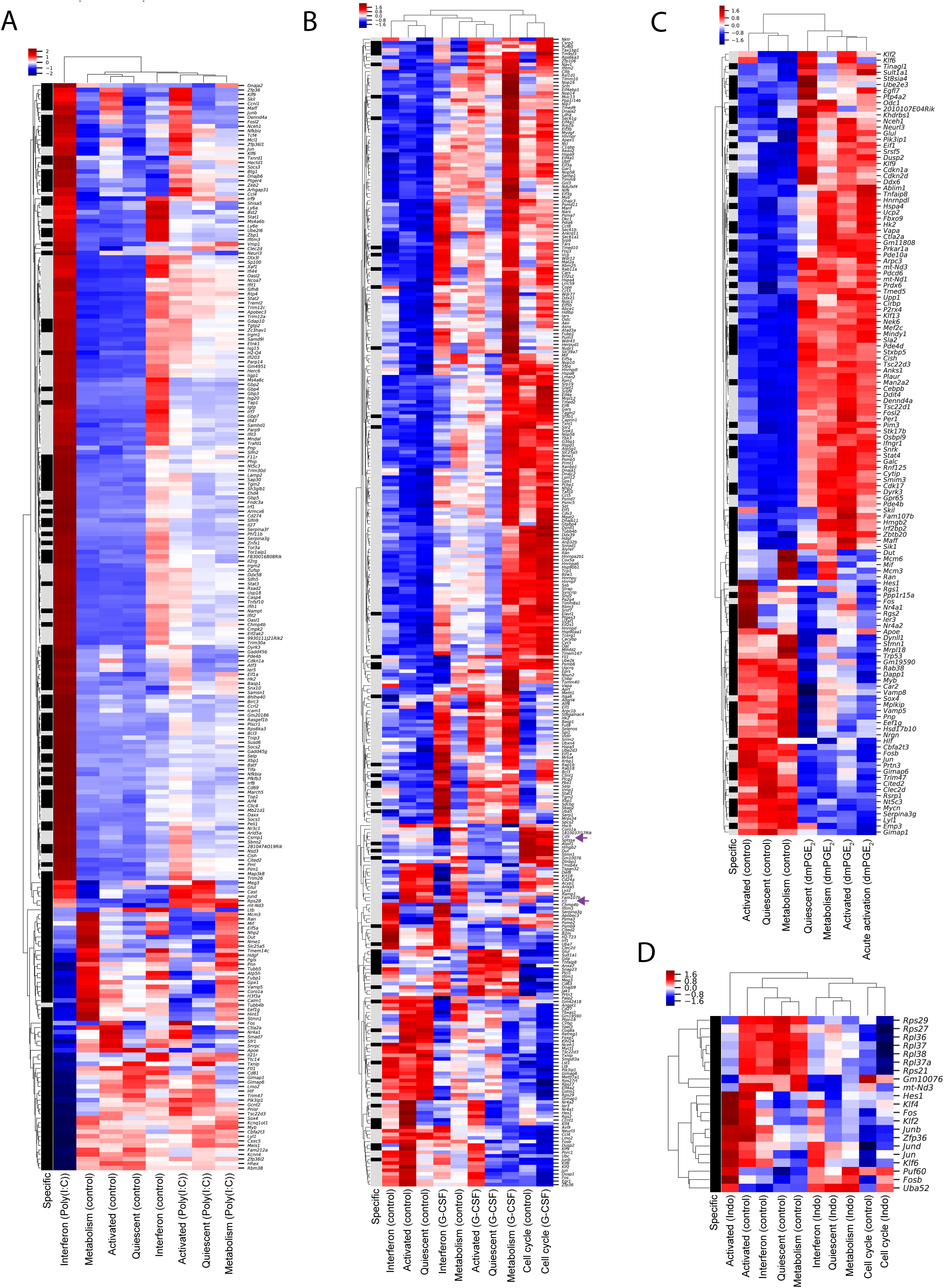
Clustered heat maps of differentially expressed genes in HSCs enables identification of genes and single cell clusters with similar expression patterns. (A-D) Heat map of differentially expressed genes between niche stimulants and control in HSCs. Single cell expression is averaged within a single cell cluster, scaled to z-scores and similar genes (rows) and clusters (columns) are aggregated by hierarchical clustering. Black row label indicates HSC specific genes, grey label marks genes differentially expressed in both HSCs and LSKs. (A) poly (I:C) at 1.5-fold cutoff. (B) G-CSF at 1.2-fold cutoff, (C) dmPGE_2_ at 1.5-fold cutoff, (D) Indomethacin at 1.2-fold cutoff.

**Supplementary Figure 6:**
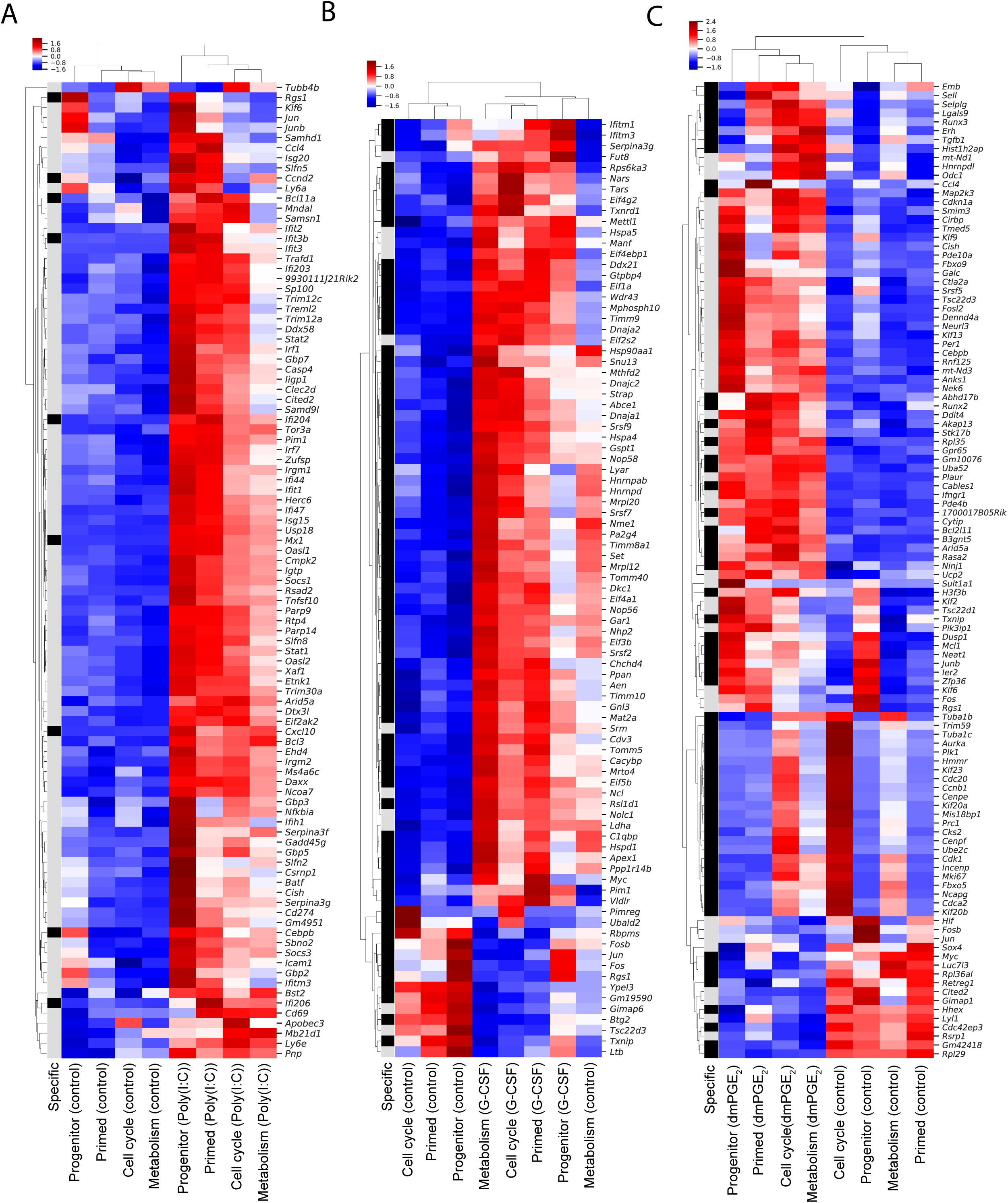
Clustered heat maps of differentially expressed genes in LSKs enables identification of genes and single cell clusters with similar expression patterns. (A-D) Heat map of differentially expressed genes between niche stimulants and control in LSKs. Single cell expression is averaged within a single cell cluster, scaled to z-scores and similar genes (rows) and clusters (columns) are aggregated by hierarchical clustering. Black row label indicates LSK specific genes, grey label marks genes differentially expressed in both HSCs and LSKs. (A) poly (I:C) at 1.5-fold cutoff. (B) G-CSF at 1.5-fold cutoff, (C) dmPGE_2_ at 1.5-fold cutoff.

**Supplemental Figure 7:**
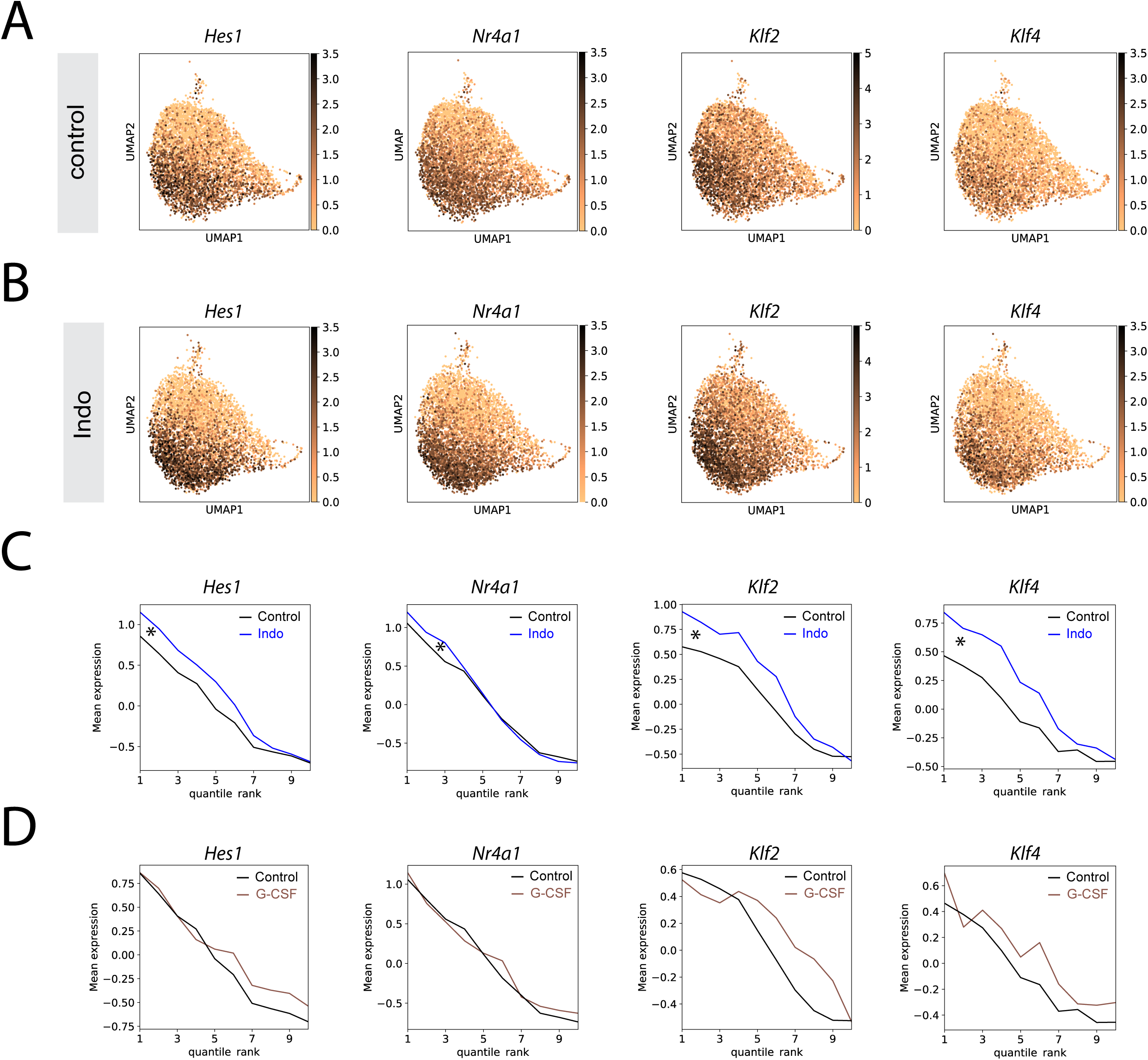
Indomethacin affects transcriptional state of IEGs (A) UMAP plot with expression of selected ‘activated’ genes in control (A) and indomethacin (B). Average expression of the same genes across cells ranked by pseudotime (cells split into 10 bins to decrease noise) comparing indomethacin and control (C, difference indicated by asterisk) or G-CSF and control (D).

**Supplementary Figure 8:**
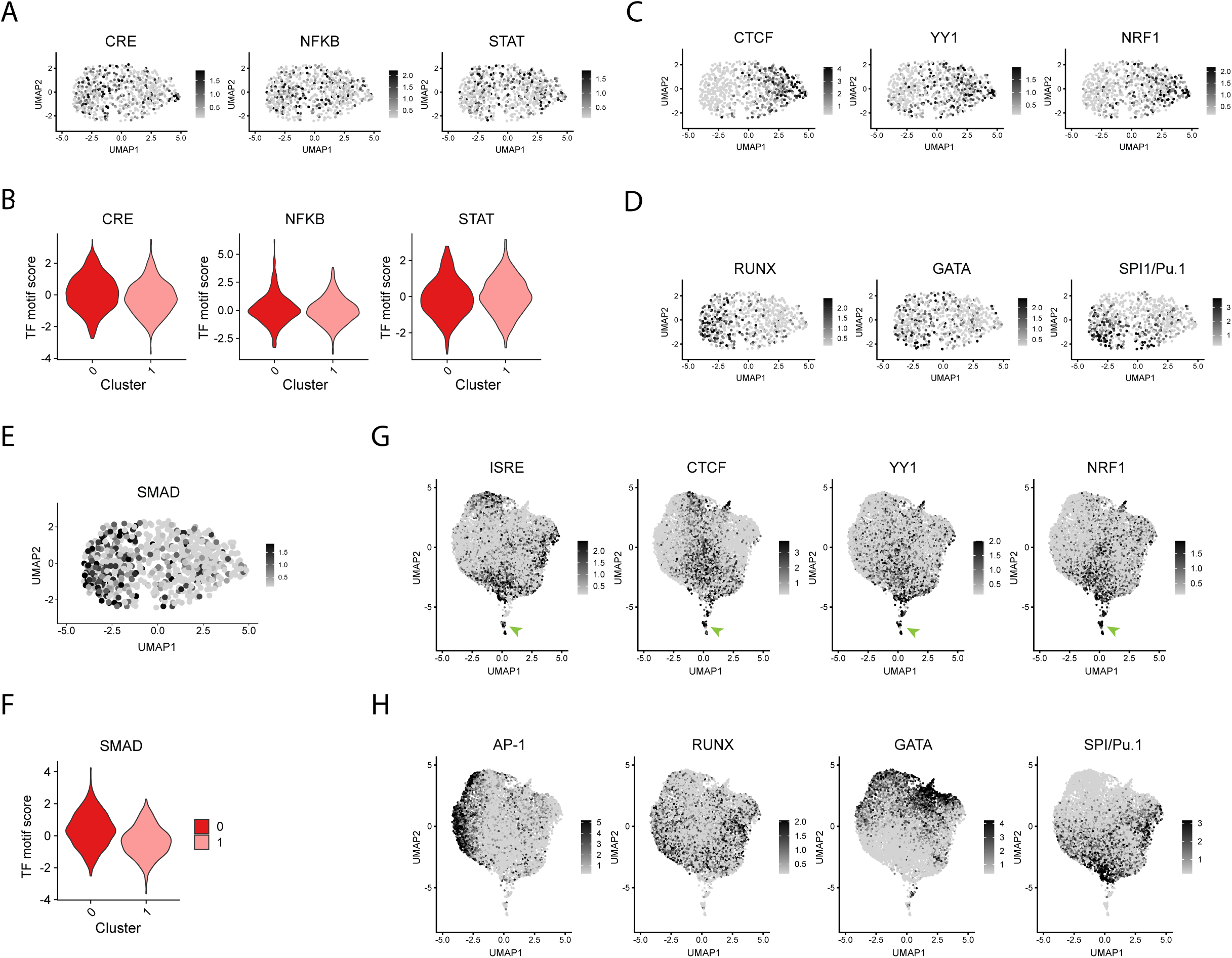
Uniform distribution of motif activity immediately downstream of niche stimulants and differential enrichment for secondary signals in HSCs and LSKs. (A-B) UMAP plots (A) and violin plots (B) of TF motif scores immediately downstream of Prostaglandin, Poly(I:C) and G-CSF signaling. (C-D) HSC UMAP plots of TF motif scores enriched in HSC cluster 1 (C) and in HSC cluster 0 (D). (E-F) UMAP plot (E) and violin plot (F) of SMAD TF motif score in HSCs. (G-H) LSK UMAP plots of TF motif scores enriched in HSC cluster 1 (G) and in HSC cluster 0 (H). Overlapping motif activities in LSK cluster 5 are indicated with a green arrowhead in G.

